# PITPβ Drives JAK2 V617F-Mediated Myeloproliferative Neoplasms by Promoting PtdIns(3,4)P_₂_-Dependent AKT Hyperactivation

**DOI:** 10.64898/2026.03.26.714558

**Authors:** Nikita A. Vantsev, Liang Zhao, Shin Morioka, Hiroaki Kajiho, Junko Sasaki, Takehiko Sasaki, Charles S. Abrams, Wei Tong

## Abstract

JAK2 is a key regulator of cytokine-mediated proliferative signaling in hematopoietic stem and progenitor cells. Activating mutations, most commonly JAK2 V617F, trigger aberrant cytokine signaling driving the pathogenesis of myeloproliferative neoplasms (MPNs). Phosphatidylinositol transfer proteins (PITPs) facilitate phosphoinositide synthesis by delivering phosphatidylinositol to lipid kinases, though their roles in oncogenic signaling have remained poorly defined. Here we show that PITPβ is critical for the development of JAK2V617F-driven MPN in mice. Deleting *Pitp*β across the hematopoietic system, but not *Pitp*α, prolonged 25-week survival of Jak2V617F mice from 10% to 85%. Loss of *Pitp*β attenuated disease-associated splenomegaly and curtailed erythroid progenitors expansion both *in vivo* and *in vitro*. Mechanistically, PITPβ is necessary for AKT hyperactivation in hematopoietic progenitors, while STAT5 and ERK signaling remain unaffected. In alignment with this role, PITPβ promotes the production of PtdIns(3,4)P_₂_, a phosphoinositide that sustains aberrant AKT signaling in Jak2V617F progenitors. Pharmacologic inhibition of AKT with the FDA-approved inhibitor capivasertib in Jak2V617F-transplanted mice similarly reduced splenomegaly and erythroid proliferation, mimicking the effects of *Pitp*β loss. Collectively, these results identify a novel PITPβ-PtdIns(3,4)P_₂_ signaling axis that selectively maintains pathological AKT activation in JAK2V617F-driven MPN, revealing a promising therapeutic vulnerability.

## INTRODUCTION

Myeloproliferative Neoplasms (MPN) are clonal hematopoietic stem cell (HSC) disorders that collectively affect approximately 300,000 people in the United States. The three principal subtypes are polycythemia vera (PV), essential thrombocythemia (ET), and primary myelofibrosis (PMF). JAK2V617F is the most common driver mutation, present in over 50% of all MPN patients and over 95% of PV patients.^1–4^ This mutation constitutively activates the JAK2 kinase, driving downstream signaling in the absence of cytokines and increasing receptor sensitivity to cytokine stimulation. Although JAK2 inhibitors help improve patient outcomes, they are not curative, suggesting the need for new therapeutic strategies.^5,6^ The mouse model that expresses Jak2 V617F (Jak2^VF^) in the endogenous locus develops a lethal MPN, characterized by elevated hematocrit, splenomegaly, and erythroid expansion in the blood, spleen, and bone marrow, mimicking PV in patients.^7^ In this study, we identify a previously unrecognized role of phosphatidylinositol transfer proteins (PITPs) in Jak2^VF^-driven MPN, whose disruption mitigated lethality and other disease manifestations in mice.

PITPs were originally identified and named after their capacity to bind and shuttle phosphatidylinositols (PtdIns) and phosphatidylcholines (PtdCho) between membrane bilayers in vitro.^8–10^ The structure of PITPs allows them to accommodate a single PtdIns or a PtdCho molecule within an internal hydrophobic pocket, which only occurs when the protein is docked at a membrane.^11–13^ Of the two class I PITP paralogs, PITPβ has a higher membrane-binding affinity and a higher phospholipid transfer rate than PITPα.^14^ Two paralogs share 77% sequence similarity but have some divergent roles in the cell. Mice lacking *Pitp*α are born with a wide range of abnormalities leading to postnatal lethality within two weeks.^15^ *Pitp*β deficiency was also initially found to cause embryonic lethality^16^, but a more recent study demonstrated no obvious defects for at least 12 months of age.^17^ Mouse embryos which are homozygous null for both *Pitp*α and *Pitp*β were not observed, suggesting that the two genes provide overlapping essential functions. There are also differences in subcellular organization between the paralogs with PITPβ primarily localizing to the cytoplasm and the trans-Golgi network and PITPα localizing to the cytoplasm and the nucleus.^18,19^

More recently, evidence emerged that PITP proteins participate in cellular signaling. The prevailing hypothesis has been that PITPs are involved in phosphoinositide synthesis by either shuttling PtdIns to the membrane sites of phosphorylation, or by interfacing with phosphatidylinositol 4-kinase (PI4K) directly to promote PtdIns phosphorylation. To this end, PITPβ has been shown to stimulate phosphoinositide synthesis in zebrafish^20^, and to be involved in PtdIns(4)P-dependent signaling during Golgi biogenesis in neuronal development in mice.^17^ We further identified an essential role for PITPs in maintaining PtdIns(4)P and PtdIns(4,5)P_2_ levels in platelets during thrombin stimulation.^21^

A growing body of evidence suggested that phosphoinositides are important for vesicular signaling in cells.^22,23^ This led us to investigate PITPs in megakaryocytes, highly secretory cells that regulate the hematopoietic system via α-granule directed release of growth factors and cytokines. Megakaryocyte and platelet-specific knockout of both *Pitp*α and *Pitp*β, via the PF4-cre, did not reduce the number of megakaryocytes in mice, but resulted in cellular structure defects and oversecretion of α-granules. This in turn resulted in over secretion of transforming growth factor β1 (TGF-β1), leading to indirect suppression of HSC and megakaryocyte progenitor proliferation. TGF-β1 blocking antibody treatment rescued PITP deficiency and restored HSC numbers.^24^

Given their role in lipid kinase signaling and their indirect effects on hematopoiesis, we hypothesized that PITPα or PITPβ might be critical mediators of signaling dysregulation in Jak2 V617F-mediated MPN. To test this hypothesis, we used Vav-cre -mediated pan-hematopoietic deletion to assess the cell-intrinsic roles of PITP isoforms in MPN. Conditional deletion of *Pitp*β, but not *Pitp*α, markedly improved survival rate, normalized red blood cell count, reduced splenomegaly, and limited erythroid progenitor expansion in Jak2^VF^ mice. Mechanistically, *Pitp*β deficiency attenuated stem cell factor (SCF) and erythropoietin (EPO)-driven AKT hyperactivation in Jak2^VF^ splenic progenitors without affecting STAT5 or ERK activation. Pitpβ is crucial for PtdIns(4,5)P_2_ generation that mediates the hyperactivation of AKT in Jak2^VF^ progenitors. We also found that the FDA-approved AKT inhibitor capivasertib reduced disease burden in Jak2^VF^-transplanted mice. Together, these findings further underscore the importance of AKT in MPN signaling and, critically, identify PITPβ as a potential therapeutic target in MPN.

## RESULTS

### *Pitp*β deficiency markedly improved poor survival and ameliorated erythrocytosis in Jak2^VF^ mice

To evaluate the effect of PITP proteins on Jak2^VF^-mediated MPN, we first generated mice with conditional knockout of either *Pitp*α or *Pitp*β using Vav-cre-mediated pan-hematopoietic deletion. We crossed either *Pitp*α^fl/fl^;Cre^vav^ or *Pitp*β^fl/fl^;Cre^vav^ into a Jak2^fl/+^ line containing an inverted V617F-mutated exon flanked by loxP sites.^7^ Henceforth, Jak2^V617F/+^;Cre^vav^ mice will be referred to as Jak2^VF^, *Pitp*α^fl/fl^;Cre^vav^ as *Pitp*α^Δ/Δ^, and *Pitp*β^fl/fl^;Cre^vav^ as *Pitp*β^Δ/Δ^. We were unable to generate adult mice with pan-hematopoietic knockout of both *Pitp*α and *Pitp*β, on either the wild-type background or Jak2^VF^ background, presumably due to embryonic lethality. Similar to earlier publications^7^, Jak2^VF^ mice developed a lethal MPN with poor survival. Only 10% of Jak2^VF^ mice survived past 25 weeks. We found that *Pitp*β deletion in pan-hematopoietic cells significantly improved survival of Jak2^VF^ mice, with over 85% of double-transgenic mice surviving past 25 weeks. *Pitp*α deletion did not have nearly the same effect, with only 33% of double-transgenic mice surviving past 25 weeks. (Figure 1A).

**Figure 1.**
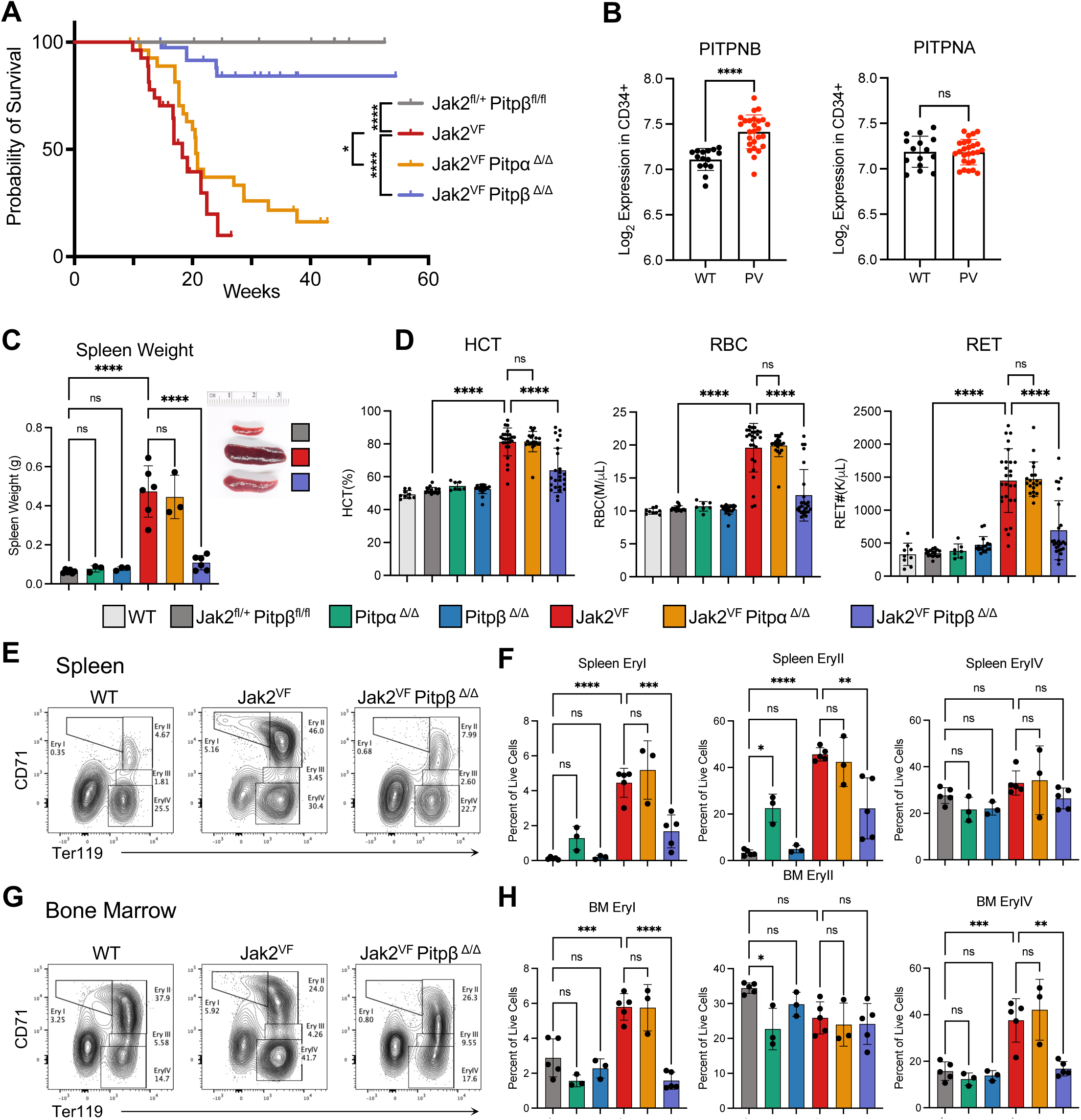
*Pitp*β deficiency improves survival and reduces erythrocytosis of Jak2^VF^ mice. (A) Event-free survival is graphed in Kaplan-Meier curves for Jak2^VF^ mice with or without combined loss of *Pitp*α or *Pitp*β. n=27-41 mice per group. P-values are calculated by log-rank analysis. (B) PITPNB (left) or PITPNA (right) mRNA levels in BM CD34+ cells^28^ from healthy controls (n=15) or JAK2^V617F+^ PV patients (n=26). (C) The weights and representative pictures of the spleens from 2- to 3-month-old mice are shown. (D) CBC analysis of peripheral blood from WT, Jak2^fl/+^;*Pitp*β^fl/fl^, *Pitp*α^Δ/Δ^, *Pitp*β^Δ/Δ^, Jak2^VF/+^;*Pitp*α^Δ/Δ^, or Jak2^VF/+^;*Pitp*β^Δ/Δ^ mice. Hematocrit: HCT; Red Blood Cell: RBC; Reticulocyte: RET. (E, F) Representative flow cytometry plot (E) and quantification (F) of erythroblasts in the spleen. (G, H) Representative flow cytometry plot (G) and quantification (H) of erythroblasts in the BM. Ery: erythroid. Ery I: CD71^high^ Ter119^-^; Ery II: CD71^high^ Ter119^+^; Ery IV: CD71^-^ Ter119^+^ population. In all relevant panels, each symbol represents an individual mouse. Bars indicate mean frequencies, and error bars indicate SD. P-values for human gene expression in CD34+ cells were calculated using Welch’s unpaired t-test. P-values for other plots were calculated using one-way ANOVA with Tukey’s multiple-comparison posttests (*, p<0.05; **, p<0.01; ***, p<0.001; ****; p<0.0001).

The striking rescue of lethality by the loss of *Pitp*β but not *Pitp*α prompted us to analyze the expression of these genes and their human orthologs in the hematopoietic system. Using publicly available genome-wide transcriptome profiling^25–27^, we found that *Pitp*β*/PITPNB* is expressed at a higher level than *Pitp*α*/PITPNA* in hematopoietic stem and progenitor cells (HSPCs) and committed progenitors in both mice and humans (Figure S1). We then analyzed the expression of *PITPNB* and *PITPNA* in human MPN. Genome-wide transcriptomic profiling of bone marrow-derived CD34+ cells^28^ from patients with JAK2^V617F+^ polycythemia vera revealed increased *PITPNB* expression compared to healthy controls, whereas *PITPNA* levels were unchanged (Figure 1B).

Analyzing mice further, we confirmed that Jak2^VF^ mice developed splenomegaly, which was alleviated by *Pitp*β deficiency. *Pitp*α deficiency did not reduce the spleen weight of Jak2^VF^ mice (Figure 1C). We then performed Complete Blood Count (CBC) analysis. As previously reported^7^, Jak2^VF^ induced an increase in hematocrit (HCT), red blood cell (RBC) count, hemoglobin amount (HGB), and reticulocyte cell count (RET), but not white blood cells (WBC) or platelets (PLT) (Figure 1E and Figure S2). *Pitp*β but not *Pitp*α deficiency significantly reduced all erythroid parameters in the peripheral blood of Jak2^VF^ mice (Figure 1E).

To further characterize the rescue of erythroid expansion, we evaluated these mice using flow cytometry. Jak2^VF^ mouse model presents with an increase in proerythroblasts (EryI: Ter119^-^CD71^+^) and basophilic/polychromatic erythroblasts (Ery II: Ter119^+^CD71^+^) in the spleen^7^, which was recapitulated in this study. *Pitp*β but not *Pitp*α deficiency significantly reduced splenic Ery I and Ery II populations in Jak2^VF^ mice (Figure 1F, G). In the bone marrow, Jak2^VF^ mice had an increased number of proerythroblasts (Ery I: Ter119^-^CD71^+^) and mature erythroblasts (Ery IV: Ter119^+^CD71^-^). Both, Ery I and Ery IV, populations were significantly reduced in Jak2^VF^;*Pitp*β^Δ/Δ^ mice, but not in Jak2^VF^;*Pitp*α^Δ/Δ^mice (Figure 1H, I).

To analyze other hematopoietic lineages, we excluded the enucleated RBC population (Ery IV, Ter119^+^CD71^-^) to more accurately quantify nucleated cells. In the spleens, while the proportion of B-cells, T-cells, and monocytes was lower in Jak2^VF^ and Jak2^VF^;*Pitp*β^Δ/Δ^ mice than that in the wild type mice, the absolute number of any of the non-erythroid cells was not significantly different (Figure S3). In Jak2^VF^ bone marrow, the proportion and the absolute cell number of B-cells were significantly reduced, which was rescued by *Pitp*β deficiency (Figure S4B,G). Other mature lymphoid and myeloid populations in the bone marrow were not affected by the Jak2^VF^ mutation or *Pitp*β deficiency (Figure S4).

### *Pitp***β** is required for Jak2^VF^ erythroid progenitor expansion of Jak2^VF^ mice

To evaluate hematopoietic stem and progenitor cells (HSPCs) and early erythroid progenitors, we combined a flow cytometry panel that identifies CFU-E and BFU-E populations^29^ with a panel that identifies multipotent progenitors (MPPs) and hematopoietic stem cells (HSCs) in mice^30^ using a spectral flow cytometer. Hematopoietic progenitors can be defined as Lineage^-^ (Lin^-^) cKit^+^ cells (LK), which are further divided into HSPC, which are Sca-1^+^ (LSK). From this population, lymphoid-biased MPP4 is LSK Flt3^+^; myeloid-biased MPP3 is LSK Flt3^-^CD48^+^CD150^-^; erythroid-biased MPP2 is LSK Flt3^-^CD48^+^CD150^+^; long-term HSCs (LT-HSC) are LSK Flt3^-^CD48^-^CD150^+^, and short-term HSCs (ST-HSC) are LSK Flt3^-^CD48^-^CD150^-^. Erythroid progenitors, or EryP, can be identified as LK CD55^+^CD105^+^CD49f^-^ cells, which is further subdivided into CFU-E (EryP CD71^hi^CD150^low^) and BFU-E (EryP CD71^low^CD150^hi^) populations (Figure 2A). Spleens of Jak2^VF^ mice exhibited a significant expansion in LK population. The expansion persisted throughout the erythroid progenitor compartments, with expansion in CD55^+^, EryP, and CFU-E/BFU-E populations, as was shown previously.^31^ HSPC population (LSK) was also elevated in the spleen of Jak2^VF^ mice. Within this population, MPP2, MPP3, ST-HSC, and LT-HSC are all elevated by the Jak2^VF^ mutation. MPP4 population could not be quantified accurately in the spleen due to low cell numbers. *Pitp*β deficiency significantly reduced Jak2^VF^-mediated expansion of erythroid progenitors, including CFU-E and BFU-E, as well as erythroid-biased MPP2, myeloid-biased MPP3, and ST-HSCs, but not LT-HSCs, in the spleen (Figure 2A, B). In the bone marrow of Jak2^VF^ mice, erythroid progenitor populations as well as MPP2 and MPP3 were elevated, similarly to that in the spleen. Jak2^VF^ mice did not exhibit an expansion of MPP4, ST-HSC, or LT-HSC populations in the bone marrow. *Pitp*β deficiency decreased all affected populations in the bone marrow (Figure 2A, C). In contrast, *Pitp*α deficiency did not reduce Jak2^VF^-mediated expansion of any of the mentioned populations (Figure 2).

**Figure 2.**
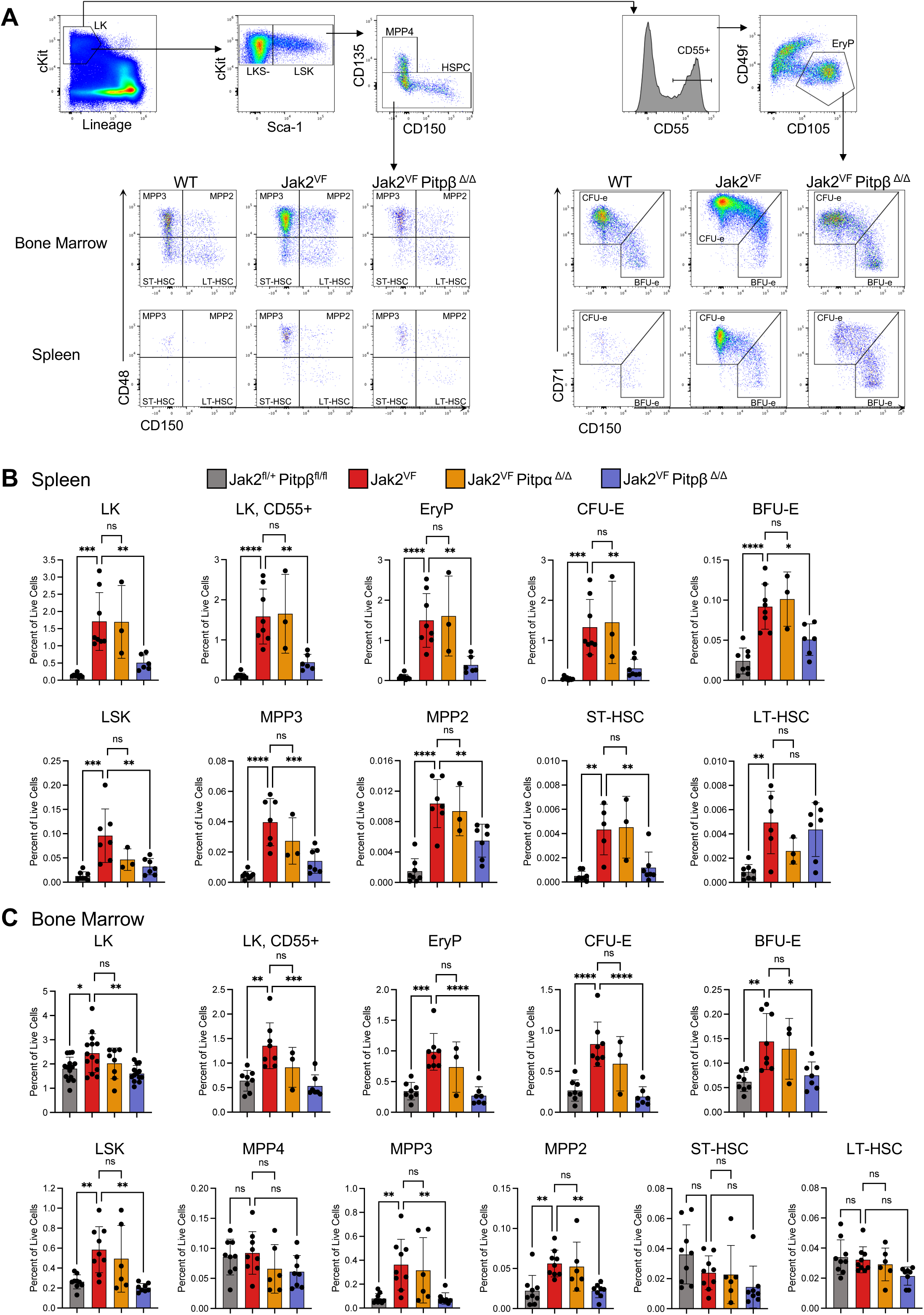
*Pitp*β deficiency reduces erythroid progenitor expansion of Jak2^VF^ mice. (A) Representative flow cytometry plots of erythroid progenitor cells and HSPCs in the BM and spleen. (B-C) Quantification of erythroid progenitor cells and HSPCs in the spleen (B) and BM (C). LK: Lin^-^cKit^+^; EryP: LK CD55^+^CD105^+^CD49f^-^; CFU-E: EryP CD71^high^CD150^low^; BFU-E: EryP CD71^low^CD150^high^; LSK: Lineage^-^cKit^+^Sca-1^+^; MPP4: LSK Flk2^+^CD150^-^; MPP3: LSK Flk2^-^CD48^+^CD150^-^; MPP2: LSK Flk2^-^CD48^+^CD150^+^; ST-HSC: LSK Flk2^-^CD48^-^CD150^-^; LT-HSC: LSK Flk2^-^CD48^-^CD150^+^. In all relevant panels, each symbol represents an individual mouse. Bars indicate mean frequencies, and error bars indicate SD. P-values were calculated using one-way ANOVA with Tukey’s multiple-comparison posttests (*, p<0.05; **, p<0.01; ***, p<0.001; ****; p<0.0001).

### *Pitp***β** is required for increased colony formation of Jak2 V617F mice

Next, we wanted to test if *Pitp*β deficiency leads to a functional change in Jak2^VF^ erythroid progenitors. Using methylcellulose colony forming assay for erythroid progenitors, we found that Jak2^VF^ cells significantly increased CFU-E colony formation compared to wild type controls even in the absence of EPO. At both low and high concentrations of EPO, Jak2^VF^ bone marrow produced more CFU-E colonies compared to that in controls. *Pitp*β deficiency reduced colony formation to normal level in all EPO concentrations tested (Figure 3 F).

**Figure 3.**
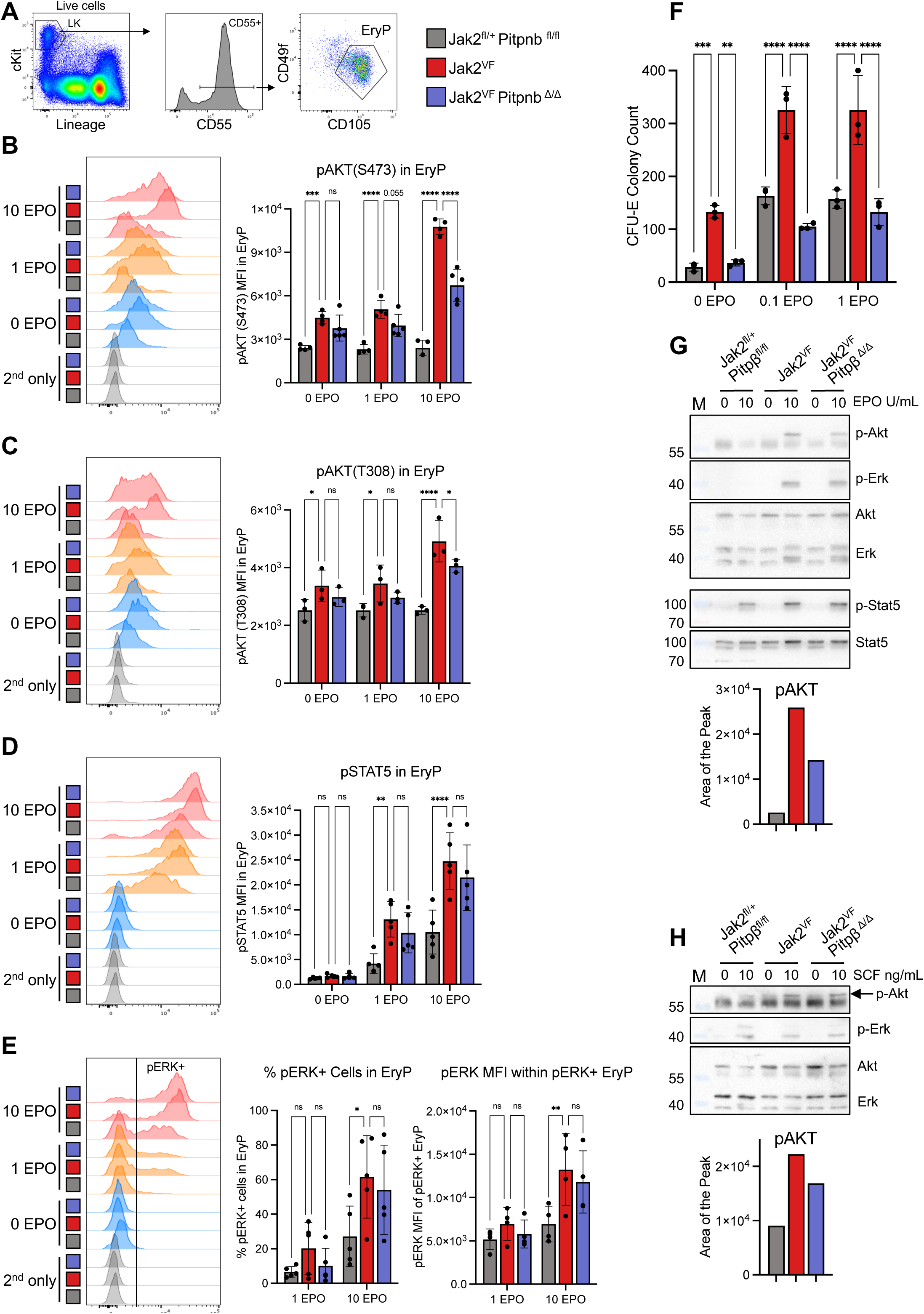
*Pitp*β deficiency reduces EPO-mediated AKT, but not STAT5 or ERK, hyperactivation in Jak2^VF^ splenic erythroid progenitors. (A) Representative flow cytometry plots and gating strategy of fixed and permeabilized splenic cells. (B-E) Representative flow cytometry plots and quantification of median fluorescence intensity (MFI) of the indicated phospho-proteins in erythroid progenitors: (B) pAKT(S473), (C) pAKT(T308), (D) pSTAT5, (E) pERK. (B-D) Total MFIs are shown; (E) pERK+ % and the MFI of pERK+ cells are shown. (F) CFU-E colony counts per 200,000 BM cells plated in semi-solid methylcellulose culture containing a graded dose of EPO in triplicate. (G) Western blot of sorted splenic EryPs stimulated with 0 or 10 U/mL EPO and pAKT quantification. (H) Western blot of sorted splenic LK cells stimulated with 0 or 10 ng/mL SCF and pAKT quantification. LK: Lin^-^cKit^+^, EryP: LK CD55^+^CD105^+^CD49f^-^. In all relevant panels, each symbol represents an individual mouse. Bars indicate mean frequencies, and error bars indicate SD. P-values were calculated using two-way ANOVA with Tukey’s multiple-comparison posttests (*, p<0.05; **, p<0.01; ***, p<0.001; ****; p<0.0001).

### *Pitp***β** is critical for EPO-induced AKT, but not STAT5 or ERK, hyperactivation in Jak2^VF^ splenic erythroid progenitors

To determine which signaling pathway is disrupted by the loss of *Pitp*β in Jak2^VF^ mice, we developed a new phospho-flow strategy to track phosphorylation of AKT, STAT5, and ERK in rare erythroid and other hematopoietic progenitors on a single-cell level. We stimulated total splenic cells with a graded dose of erythropoietin (EPO) and measured phosphorylation of signaling proteins in different erythroid progenitor and HSPC populations (Figure 3A, Figure S5A). We observed that while many, but not all, cells within the Ery I population had a marked increase in pAKT and pSTAT5 median fluorescence intensity (MFI) in response to EPO. In comparison, more mature Ery A or Ery B populations did not respond to EPO (Figure S5C). The LK population showed a heterogeneous response to EPO stimulation, as expected. CD55 specifically subdivided LK cells into EPO-responding (CD55^+^) and non-responding (CD55^-^) cells (Figure S5C, D), in agreement with prior studies that identify CD55 as an indicator of erythroid/meg commitment.^29,32^ Within the LK CD55^+^ population, EryP (LK CD55^+^CD105^+^CD49f^-^) cells, which consist of both BFU-E and CFU-E progenitors, had the strongest response to EPO. This was consistent between EPO-stimulated pAKT or pSTAT5 activation (Figure S5 B-F). We concluded that EryP is more enriched in erythroid progenitor cells compared to Ery I. The subsequent studies were focused on EryP.

Upon EPO stimulation, splenic EryPs from Jak2^VF^ mice displayed a significantly higher activation of pAKT (S473), pAKT (T308), pSTAT5, and pERK compared to WT counterparts, as indicated by MFI. Importantly, *Pitp*β deficiency reduced EPO-stimulated AKT phosphorylation at both S473 and T308 but did not significantly reduce phosphorylation of STAT5 or ERK in Jak2^VF^ mice (Figure 3B-E). While the bone marrow EryP population showed a robust response to EPO, Jak2^VF^ mice did not exhibit an increase in signaling compared to control mice, thus we focused on splenic progenitors in subsequent studies.

To confirm our findings, we sorted EryP population from spleens of WT, Jak2^VF^, and Jak2^VF^;*Pitp*β^Δ/Δ^ mice, stimulated them with EPO and measured activation of phospho proteins by western blot (WB). Similar to phosphoflow, splenic EryP from Jak2^VF^ mice exhibited hyperactivated pAKT, pSTAT5, and pERK in response to EPO compared to the controls. The increase in pAKT was partially reduced by *Pitp*β deficiency (Figure 3G). This suggests that Pitpβ does not directly affect JAK2/STAT5 or ERK signaling, but rather specifically impacts AKT activation.

### *Pitp***β** is critical for SCF-mediated AKT hyperactivation in splenic myeloid progenitors and HSPCs of Jak2^VF^ mice

Next, we analyzed whether *Pitp*β deficiency reduced AKT hyperactivation in response to cytokines other than EPO. To test this, we employed a similar phosphoflow strategy that focuses on HSPCs and myeloid progenitors. We stimulated splenocytes with stem cell factor (SCF) and evaluated pAKT activation. We found that progenitor cells (LK) and HSPCs (LSK) from Jak2^VF^ mice have significantly elevated AKT phosphorylation in response to SCF compared to WT. Jak2^VF^ also increased pAKT in MkP (megakaryocyte progenitors, LKS^-^CD41^+^CD150^+^), and GMP (granulocyte-monocyte progenitors, LKS^-^CD41^-^FcγRII/III^+^) populations in response to SCF. *Pitp*β deficiency significantly reduced SCF-stimulated pAKT in LK, LSK, MkP, and GMP hyperactivated by Jak2^VF^ (Figure 4 B). SCF did not stimulate STAT5 phosphorylation in any genotype (Figure S6B), as expected, again highlighting the specific role of Pitpβ in AKT signaling. We confirmed the phosphoflow data by WB. Sorted LK cells were stimulated with SCF. Jak2^VF^ splenic LK progenitors showed hyperactivated pAKT, which was reduced upon the loss of *Pitp*β (Figure 3H).

**Figure 4.**
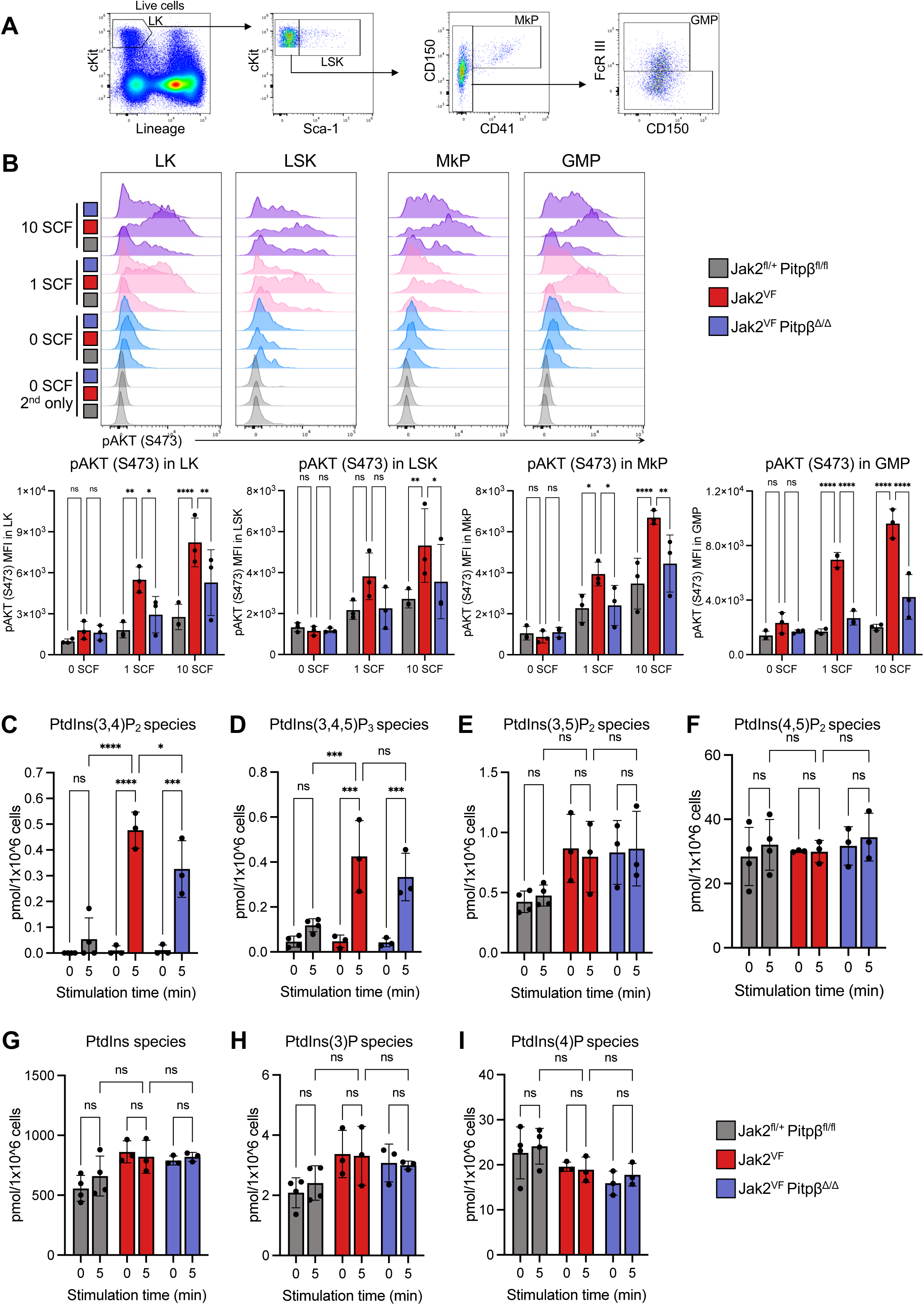
*Pitp*β deficiency reduces SCF-mediated PtdIns(3,4)P_2_ generation and subsequent AKT hyperactivation in Jak2^VF^ splenic progenitors. (A) Representative flow cytometry plots and gating strategy of fixed and permeabilized splenic cells. (B) Representative flow cytometry plots and quantification of median fluorescence intensity (MFI) of pAKT(S473) of indicated progenitor populations. LK: Lin^-^cKit^+^; LSK: Lin^-^cKit^+^Sca-1^+^; LKS^-^: Lin^-^cKit^+^Sca-1^-^; MkP: LKS^-^ CD150^+^CD41^+^; GMP: LKS^-^ CD41^-^ FcγRIII^+^. (C-I) LS-MS measurement of phosphoinositide species in sorted LK population treated with 0 or 10ng/mL SCF for 5 min. Total sum of molecular species is shown for each phosphoinositide. PtdIns(5)P was not detected. In all relevant panels each symbol represents an independent biological replicate containing 1-4 mice each. Bars indicate mean frequencies, and error bars indicate SD. P-values were calculated using two-way ANOVA with Tukey’s multiple-comparison posttests (*, p<0.05; **, p<0.01; ***, p<0.001; ****; p<0.0001).

### *Pitp***β** is critical for PtdIns(3,4)P_2_ generation, mediating AKT hyperactivation in Jak2^VF^ progenitors

We previously showed that PITPs are required for PtdIns(4)P and PtdIns(4,5)P_2_ production during thrombin stimulation in platelets.^21^ Since PITPβ is required for AKT hyperactivation in Jak2^VF^ splenic hematopoietic progenitors, we hypothesized that it would also be required for the cytokine stimulated production of PtdIns(3,4,5)P_3_ and/or PtdIns(3,4)P_2_, two type of phosphoinosities important for PI3K-AKT activation.^33, 34^ To test this hypothesis, sorted splenic LK cells were stimulated with SCF for 5 min, and phospholipids were analyzed by Mass Spectrometry. SCF stimulated an increase in PtdIns(3,4,5)P_3_ and PtdIns(3,4)P_2_ levels, but not phosphatidylinositol or other species of phosphoinositides, such as PtdIns(3)P, PtdIns(4)P, PtdIns(3,5)P, or PtdIns(4,5)P_2_, in mice from all genotypes (Figure 4C-I). Upon SCF stimulation, splenic progenitors from Jak2^VF^ mice exhibited a significantly higher level of PtdIns(3,4,5)P_3_ and PtdIns(3,4)P_2_ compared to WT control (Figure 4C, D). Importantly, *Pitp*β deficiency significantly reduced PtdIns(3,4)P_2_ level in Jak2^VF^ mice (Figure 4C), and showed a downward trend for PtdIns(3,4,5)P_3_ level, albeit not statistically significant (Figure 4D).

### AKT inhibitor reduced PV phenotype in JAK2^VF^ transplanted mice

Our data suggest that *Pitp*β deficiency rescues Jak2^VF^ MPN phenotype by specifically reducing AKT signaling. Previous studies demonstrated that pharmacologic AKT inhibition with MK-2206 reduces Jak2^VF^-driven disease burden in mice.^35,36^ Building upon this observation, we tested capivasertib (AZD-5363), a recently FDA-approved orally administered drug against breast cancer.^37,38^ We transplanted Jak2^VF^ bone marrow cells into wild type recipients and confirmed development of disease, as evidenced by elevated HCT, RBC, and RET counts by CBC analysis 3 weeks after transplantation. Mice were then randomized to receive either Capivasertib (150mg/kg b.i.d.) or vehicle for 4 weeks (Figure 5A). We observed a reduction in hematocrit, red blood cells and reticulocyte counts during this treatment period (Figure 5B). Following completion of the 4-week treatment period, we found a reduction in the spleen weight, spleen cell count, and bone marrow cell count in transplanted mice treated with Capivasertib (Figure 5C, D). Flow cytometry analysis showed a significant decrease in the number of proerythroblasts (Ery I) in the bone marrow and spleens of capivasertib-treated mice (Figure 5E). Moreover, capivasertib reduced LK, LK CD55^+^, EryP, and CFU-E/BFU-E populations in the spleen (Figure 5F) and the bone marrow (Figure 5G). Taken together, our findings strengthen the evidence that AKT inhibition, now achievable with an FDA approved drug, can alleviate Jak2^VF^-associated symptoms.

**Figure 5.**
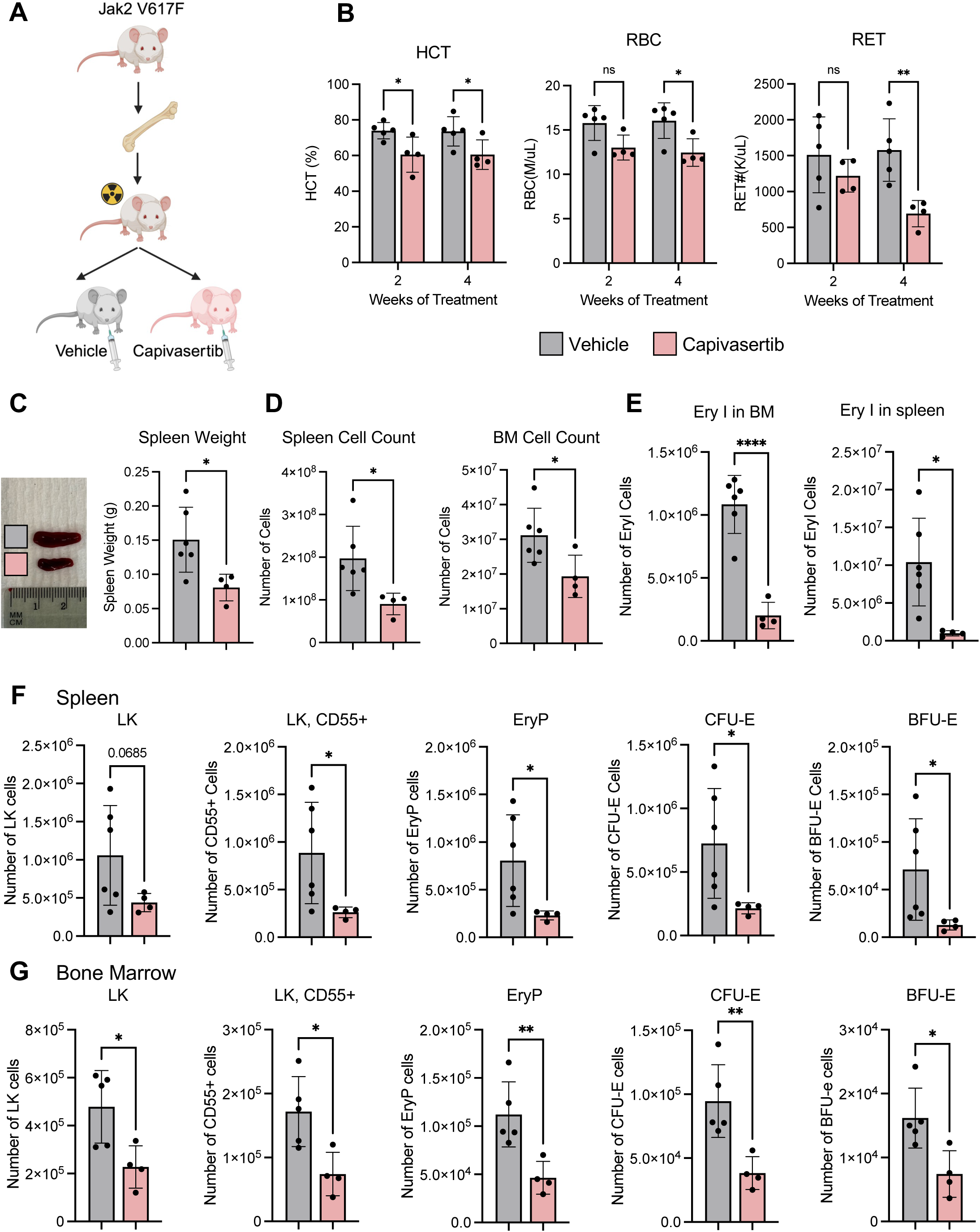
AKT inhibitor reduces Jak2^VF^-mediated erythroid expansion in mice. (A) Schematic illustration of capivasertib treatment of mice transplanted with Jak2^VF^ BM. (B) CBC analysis of Jak2^VF^ transplanted mice 2 or 4 weeks after capivasertib or vehicle treatment. Hematocrit: HCT; Red Blood Cell: RBC; Reticulocyte: RET. (C) Representative picture (left) and weights (right) of spleens from mice 4 weeks after Capivasertib or vehicle treatment. (D) Spleen (left) and BM (right) cell counts of mice after 4 weeks of treatment. (E-G) Flow cytometry analysis of erythroid populations in treated mice, (E) Quantification of the number of Ery I erythroid progenitors (CD71^+^ Ter119^-^) in the BM (left) and spleen (right). (F-G) Quantification of the number of CFU-E and BFU-E progenitors in the spleen (F) and BM (G). LK: Lin^-^ cKit^+^; EryP: LK CD55^+^CD105^+^CD49f^-^; CFU-E: EryP CD71^high^CD150^low^; BFU-E: EryP CD71^low^ CD150^high^. In all relevant panels, each symbol represents an individual mouse. Bars indicate mean frequencies, and error bars indicate SD. P-values for the CBC analysis were calculated using two-way ANOVA with Holm-Šídák multiple-comparison posttests. P-values for other analyses were calculated using Welch’s unpaired t-test (*, p<0.05; **, p<0.01; ***, p<0.001; ****; p<0.0001).

## DISCUSSION

This work provides the first evidence that phosphatidylinositol transfer protein β is essential for MPN development in mice harboring the endogenous Jak2 V671F mutation. *Pitp*β deletion rescues the lethality of Jak2^VF^ mutation in mice and mitigates erythrocytosis. *Pitp*β is required for the expansion of erythroid progenitors and multipotent progenitors driven by Jak2^VF^. Using our new phosphoflow strategy, we showed that Jak2^VF^ potentiates and promotes AKT, STAT5, and ERK activation by EPO in erythroid progenitors. Strikingly, Jak2^VF^ strongly enhanced AKT activation by SCF in myeloid progenitors and HSPCs. SCF signals through its receptor tyrosine kinase KIT without activating JAK2-STAT5, implying a crosstalk between KIT and type 1 cytokine receptors such as EPOR. This crosstalk has been reported^39,40^, but the mechanism remains incompletely understood. Our results suggest a specific role of Pitpβ in AKT signaling, but not JAK-STAT signaling. It is notable that pharmacologic AKT inhibition is sufficient to attenuate Jak2^VF^-mediated MPN. These findings establish Pitpβ as a potential therapeutic target in MPN and further emphasize the central role of AKT in MPN progression.

Cytokine signaling elicited a robust signaling response in progenitor cells from both bone marrow and spleen. In the spleen, Jak2^VF^ markedly hyperactivates AKT, STAT5, and ERK compared to wild type mice, as reported previously on primary mouse cells.^7,41–44^ In contrast, STAT5 hyperactivation in primary mouse bone marrow cells is less pronounced.^45^ Similarly, we also did not detect a clear hyperactivation of AKT, STAT5, or ERK in Jak2^VF^ bone marrow. It is possible that the MPN manifestations associated with Jak2^VF^ mice may be driven predominantly by an expanded pool of hyperactive erythroid progenitors specifically in the spleen. This may account for the more pronounced signaling phenotypes observed in the spleen compared to the bone marrow.

Among the two paralogs, only Pitpβ deficiency rescues Jak2^VF^ phenotype. This selective effect may reflect differences in their expression patterns (Pitpβ is more highly expressed in progenitors^24^) and/or distinct subcellular localization.^18,19^ Dissecting these features will be an important focus of future studies. Two splice variants of Pitpβ have been described, one of which contains a C-terminal serine residue that is constitutively phosphorylated by protein kinase C (PKC). Although this phosphorylation was initially proposed to be required for Golgi localization^46^, subsequent studies showed it was dispensable for this function.^47^ Its role in JAK2-driven signaling, if any, remains unresolved and merits further investigation. It is notable that *Pitp*β deficiency only partially attenuated AKT activation yet produced a striking benefit in Jak2^VF^ mouse survival and curtailed erythroid progenitor expansion. This suggests that Pitpβ may regulate additional pathways beyond AKT and underscores the need for further signaling analysis to define the full contribution of Pitpβ in Jak2^VF^-mediated MPN.

Decreased expression of Pitpα has been reported to enhance phospho-AKT signaling in muscle, which may have contributed to ameliorating dystrophic changes in Duchenne muscular dystrophy in dogs and zebrafish, suggesting context-dependent roles for PITP paralogs in AKT regulation.^48^ This could be explained by the distinct, non-equivalent function of Pitpα or Pitpβ, or a compensatory upregulation of Pitpβ upon Pitpα loss that increases lipid second messengers within the PI3K-AKT axis. More broadly, these data support a role for PITP proteins as modulators of AKT signaling.

It has been suggested that PtdIns(3,4,5)P_3_ acutely actives AKT, while PtdIns(3,4)P_2_ is required for sustained activation.^34^ Our data indicate that Jak2^VF^ induces an increase in both species and that *Pitp*β deficiency reduces PtdIns(3,4)P_2_ level. We and others have also shown that PITP proteins facilitate PI4K-dependent synthesis of PtdIns(4)P^20,21^, a precursor to PtdIns(3,4,5)P_3_. On this basis, our working model is that loss of PITPβ diminishes signaling flux through the Jak2^VF^-AKT pathway by limiting the availability of PtdIns(4)P and consequently PtdIns(3,4)P_2_ and PtdIns(3,4,5)P_3_ generation. Mass spectrometry analysis failed to detect any significant changes in PtdIns(3,4,5)P_3_ levels following 5 minutes of SCF stimulation in Jak2^VF^;*Pitp*β^Δ/Δ^ mice in comparison to Jak2^VF^ mice. PtdIns(3,4,5)P_3_ generation kinetics is fast and transient, which may contribute to the difficulty in quantifying PtdIns(3,4,5)P_3_ precisely to capture subtle changes impacted by Pitpβ loss. Furthermore, the limitation of our approach is that it quantifies total cellular phosphoinositides rather than localized pools. It is possible that Pitpβ regulates PtdIns(3,4,5)P_3_ generation in a key microdomain critical for Akt activation, which warrants future investigation.

These findings further support AKT as a rationale therapeutic strategy in MPN. Previous studies on PI3K pathway inhibitors have demonstrated clinical activity^49^, but been constrained by tolerability issues. The allosteric AKT inhibitor MK-2206 has demonstrated efficacy in preclinical MPN models, reinforcing the importance of this pathway in disease maintenance.^35,36^ In addition, the AKT inhibitor capivasertib has been proven safe and effective in patients with solid tumors, including breast cancer.^37,38^ Here, we show that capivasertib also attenuated MPN disease burden in mice, highlighting it as a promising candidate drug for therapeutic evaluation in MPN.

Beyond AKT inhibition, this work strongly supports the development of selective PITPβ inhibitors as a novel therapeutic strategy. It is notable that *Pitp*β deletion does not completely abolish AKT phosphorylation, yet still results in a marked survival benefit in mice, suggesting that partial attenuation of this pathway may offer improved tolerability compared with complete AKT blockade. Finally, elevated *PITPNB* expression in MPN patient samples further implicates PITPβ in disease progression and underscores its potential as a therapeutic target.

## METHODS

### Mice

Animal protocols were approved by the institutional animal use committees of the University of Pennsylvania. To generate transgenic mice, either *Pitp*α^fl/fl^;Cre^vav^ or *Pitp*β^fl/fl^;Cre^vav^ mice with C57BL/6 background were crossed into a Jak2^fl/+^ line containing an inverted V617F-mutated exon flanked by loxP sites.^7^ All mice were bread in-house in the animal facility at the University of Pennsylvania and maintained on standard chow. Mice of both sexes were used for the study, unless otherwise indicated, and were age matched at 2-3 months.

### CBC assay

Peripheral blood was collected into EDTA-coated tube and analyzed using IDEXX ProCyte Dx CBC hematology analyzer.

### Flow cytometry of fresh cells

Bone marrow from one femur and tibia each was spun out into PBS containing 2% Bovine Calf Serum (BCS). Spleens were harvested and shredded into single cell suspension in PBS with 2% BCS. A portion of cells was directly stained for Lineage Panel antibodies for 30 min at 4^0^C. A portion of cells was incubated with RBC lysis buffer (0.8% NH_4_Cl, 10uM EDTA, pH 7.4∼7.6) for 1 min at 4^0^C, washed, and stained with biotin-conjugated anti-lineage antibodies for 30 min at 4^0^C. Following which cells were stained with HSPC Panel antibodies for 30 min at 4^0^C. Data was acquired on Cytek Aurora cytometer and analyzed using BD FlowJo.

### Phospho flow cytometry

BM and spleen single cells were collected as described above, RBC lysed and stained with Zombie Aqua for 10 min at room temperature. Following which cells were starved and stained with either Pflow Erythroid Panel or Pflow Myeloid Panel antibodies in 250uL IMDM with 0.5% BCS and 50uL BD Horizon Brilliant Stain Buffer for 30 min at 37^0^C. Cells were then incubated with 10U/mL EPO or 10ng/mL SCF for 15 min at 37^0^C and immediately fixed with 1.5% paraformaldehyde at room temperature for 10 minutes. Cells were washed and permeabilized with ice-cold methanol at 4^0^C for 10 min and either stained immediately or stored in -80^0^C. Following permeabilization, cells were washed and incubated with primary phospho antibodies for 30 min at 4^0^C, followed by the secondary antibody for 30 min at 4^0^C. Data was acquired on Cytek Aurora cytometer and analyzed using BD FlowJo.

### Western Blot

BM and spleen single cells were collected as described above, RBC lysed and stained with Direct Lineage Cell Depletion beads for 20 min at 4^0^C. Lineage positive cells were depleted using autoMACS Pro separator. Lineage negative fraction was stained with Sorter Panel antibodies for 30 min at 4^0^C. Cells were sorted on BD Aria Fusion. Purified cells were starved in 500uL IMDM with 0.5% BCS for 30 min at 37^0^C. Cells were incubated with 10U/mL EPO or 10ng/mL SCF for 15 min at 37^0^C, then pelleted. The supernatant was aspirated, the pellet frozen on dry ice and stored in -80^0^C. Cell pellets were lysed in the following buffer: 1% NP-40, 50mM Tris, 250mM NaCl, 5mM EDTA, 3mM NaF, 3mM Na_2_VO_3_, 1mM Phenylmethylsulfonyl fluoride, 1:50 cOmplete Protease Inhibitor Cocktail.

### CFU assay

BM was collected as described above. 200,000 cells were plated per 1mL of MethoCult M3234 supplemented with 0, 0.1U/mL, or 1U/mL EPO and cultured at 37^0^C in 5% CO_2_ incubator. CFU-E colonies were enumerated 48 hours after plating.

### Bone marrow transplantation and *in vivo* Capivasertib treatment

BM from Jak2^VF^ mice (CD45.2^+^) was isolated. 2*10^6^ cells were injected into lethally irradiated (5Gy + 5Gy) female age matched WT recipient (CD45.1^+^) mice. Following a 3-week reconstitution, mice were randomly selected to receive either vehicle or capivasertib. Capivasertib oral solution was prepared in a 5% (v/v) DMSO, 25% (w/v) Kleptose HPB in water and dosed at 150mg/kg BID for 4 weeks by oral gavage.

### Cell preparation, lipid extraction, and derivatization for phosphoinositide analyses

Spleen single cells were collected as described above. The LK population was sorted as described above. Cells were starved in 500 μL IMDM containing 0.5% BCS for 30 min at 37°C. Cells were then incubated with 10 ng/mL SCF for 5 min at 37°C and immediately fixed by adding an equal volume (1:1, v/v) of 2 M HCl. The cells were then centrifuged at 2,000 x g for 5 min. The supernatant was aspirated, the pellet was frozen on dry ice and stored in −80°C. The frozen cell pellets were then subjected to phosphoinositide analysis as described previously^21,50,51^, with minor modifications.

Briefly, cell pellets were mixed thoroughly with 500 μL of formic acid and 4.5 mL of chloroform:methanol (1:1, v/v). A 3.5 mL aliquot was transferred to a new glass tube for PI(3,4,5)P_₃_ measurement, and a 1.5 mL aliquot was transferred to a separate glass tube for phosphoinositide regioisomer measurement by chiral column chromatography and mass spectrometry (PRMC-MS) analysis. For PI(3,4,5)P_₃_ measurement, 50 μL of chloroform:methanol (1:9, v/v) containing 10 pmol of C16:0/C16:0-d_₆₂_ PI(3,4,5)P_₃_ as an internal standard and 1 nmol of C8:0/C8:0 PI(4,5)P_₂_ as an adsorption protectant was added. For PRMC-MS analysis, 50 μL of chloroform:methanol (1:9, v/v) containing 10 pmol of C17:0/C20:4 phosphoinositides (PI(3,4)P_₂_, PI(3,5)P_₂_, PI(4,5)P_₂_, PI3P, PI4P, and PI5P) and 20 pmol of C15:0/C18:1-d_7_ PI as internal standards, together with 1 nmol of C8:0/C8:0 PI(4,5)P_₂_ as an adsorption protectant, was added. Concentration and derivatization of phosphoinositides were performed as described previously^47^, followed by drying under a stream of nitrogen gas. The dried samples were reconstituted in 60 μL of methanol:ultrapure water (3:1, v/v) for PI(3,4,5)P_₃_ measurement or in 60 μL of acetonitrile for PRMC-MS analysis.

### Mass spectrometry

PI(3,4,5)P_3_ measurement was performed as described previously^52^, with minor modifications. LC-MS/MS analysis was conducted in the positive-ion mode using an Ultimate 3000 LC system (Thermo Fisher Scientific) coupled to a TSQ Vantage triple-stage quadrupole mass spectrometer (Thermo Fisher Scientific). A 20 μL aliquot of the sample was injected and separated on a COSMOCORE 2.6C_18_ column (2.1 × 150 mm, 2.6 μm; Nacalai Tesque) at 60°C using mobile phase A consisting of isopropanol:acetonitrile:70% ethylamine (800:200:0.65, v/v/v) and mobile phase B consisting of acetonitrile:ultrapure water:70% ethylamine (800:200:0.65, v/v/v). The gradient program was as follows: 0–1 min, 30% A; 1–3 min, 30–90% A; 3–7.5 min, 90% A; and 7.5–13 min, 30% A, at a flow rate of 220 μL/min.

PRMC-MS was performed as described previously^51^, with minor modifications. A Nexera XR HPLC/autosampler system (Shimadzu) coupled to a triple quadrupole mass spectrometer QTRAP 6500 (SCIEX) was used. Twenty microliters of the sample was injected and separated on a CHIRALPAK IC-3 column (cellulose tris[3,5-dichlorophenylcarbamate], 2.1 × 250 mm, 3 μm; DAICEL) at 23°C using mobile phase A consisting of methanol/5 mM ammonium acetate and mobile phase B consisting of acetonitrile/5 mM ammonium acetate. The gradient program was as follows: 0–1 min, 35% A; 1–2 min, 35–70% A; 2–12 min, 70% A; and 12–20 min, 35% A, at a flow rate of 100 μL/min.

## ACKNOWLEDGEMENTS

This work was supported by T32 HL007971-22 to NV. WT is supported by NIH grants R01 CA271523 and R01CA282668. CA is supported by NIH grants R01 HL146722 and P01 HL146373. We are grateful to Siera A Tomishima for help on the phosphoflow protocol.

## AUTHOR CONTRIBUTIONS

W. T. and C. A. conceived the project and supervised the studies. L. Z. bred the mice and performed the CBC analysis. S.M., H.K., J.S., T.S. performed phospholipid LS-MS experiments. N.V. performed CBC, AKT inhibitor, and flow cytometry experiments and wrote the manuscript.

## COMPETING INTERESTS DISCLOSURES

The authors declare no competing interests.

## Notes

### Competing Interest Statement

The authors have declared no competing interest.

## REFERENCES

1 Baxter, E. J. et al. Acquired mutation of the tyrosine kinase JAK2 in human myeloproliferative disorders. Lancet 365, 1054–1061 (2005). 10.1016/S0140-6736(05)71142-9

2 James, C. et al. A unique clonal JAK2 mutation leading to constitutive signalling causes polycythaemia vera. Nature 434, 1144–1148 (2005). 10.1038/nature03546

3 Kralovics, R. et al. A gain-of-function mutation of JAK2 in myeloproliferative disorders. N Engl J Med 352, 1779–1790 (2005). 10.1056/NEJMoa051113

4 Levine, R. L. et al. Activating mutation in the tyrosine kinase JAK2 in polycythemia vera, essential thrombocythemia, and myeloid metaplasia with myelofibrosis. Cancer Cell 7, 387–397 (2005). 10.1016/j.ccr.2005.03.023

5 Harrison, C. et al. JAK inhibition with ruxolitinib versus best available therapy for myelofibrosis. N Engl J Med 366, 787–798 (2012). 10.1056/NEJMoa1110556

6 Cervantes, F. et al. Three-year efficacy, safety, and survival findings from COMFORT-II, a phase 3 study comparing ruxolitinib with best available therapy for myelofibrosis. Blood 122, 4047–4053 (2013). 10.1182/blood-2013-02-485888

7 Mullally, A. et al. Physiological Jak2V617F expression causes a lethal myeloproliferative neoplasm with differential effects on hematopoietic stem and progenitor cells. Cancer Cell 17, 584–596 (2010). 10.1016/j.ccr.2010.05.015

8 Wirtz, K. W. & Zilversmit, D. B. Participation of soluble liver proteins in the exchange of membrane phospholipids. Biochim Biophys Acta 193, 105–116 (1969). 10.1016/0005-2736(69)90063-7

9 Helmkamp, G. M., Jr., Harvey, M. S., Wirtz, K. W. & Van Deenen, L. L. Phospholipid exchange between membranes. Purification of bovine brain proteins that preferentially catalyze the transfer of phosphatidylinositol. J Biol Chem 249, 6382–6389 (1974).

10 Carvou, N. et al. Phosphatidylinositol- and phosphatidylcholine-transfer activity of PITPbeta is essential for COPI-mediated retrograde transport from the Golgi to the endoplasmic reticulum. J Cell Sci 123, 1262–1273 (2010). 10.1242/jcs.061986

11 Shadan, S. et al. Dynamics of lipid transfer by phosphatidylinositol transfer proteins in cells. Traffic 9, 1743–1756 (2008). 10.1111/j.1600-0854.2008.00794.x

12 Tilley, S. J. et al. Structure-function analysis of human [corrected] phosphatidylinositol transfer protein alpha bound to phosphatidylinositol. Structure 12, 317–326 (2004). 10.1016/j.str.2004.01.013

13 Vordtriede, P. B., Doan, C. N., Tremblay, J. M., Helmkamp, G. M., Jr. & Yoder, M. D. Structure of PITPbeta in complex with phosphatidylcholine: comparison of structure and lipid transfer to other PITP isoforms. Biochemistry 44, 14760–14771 (2005). 10.1021/bi051191r

14 Baptist, M., Panagabko, C., Cockcroft, S. & Atkinson, J. Ligand and membrane-binding behavior of the phosphatidylinositol transfer proteins PITPalpha and PITPbeta. Biochem Cell Biol 94, 528–533 (2016). 10.1139/bcb-2015-0152

15 Alb, J. G., Jr., et al. Mice lacking phosphatidylinositol transfer protein-alpha exhibit spinocerebellar degeneration, intestinal and hepatic steatosis, and hypoglycemia. J Biol Chem 278, 33501–33518 (2003). 10.1074/jbc.M303591200

16 Alb, J. G., Jr., et al. Genetic ablation of phosphatidylinositol transfer protein function in murine embryonic stem cells. Mol Biol Cell 13, 739–754 (2002). 10.1091/mbc.01-09-0457

17 Xie, Z., Hur, S. K., Zhao, L., Abrams, C. S. & Bankaitis, V. A. A Golgi Lipid Signaling Pathway Controls Apical Golgi Distribution and Cell Polarity during Neurogenesis. Dev Cell 44, 725–740 e724 (2018). 10.1016/j.devcel.2018.02.025

18 Phillips, S. E., Ile, K. E., Boukhelifa, M., Huijbregts, R. P. & Bankaitis, V. A. Specific and nonspecific membrane-binding determinants cooperate in targeting phosphatidylinositol transfer protein beta-isoform to the mammalian trans-Golgi network. Mol Biol Cell 17, 2498–2512 (2006). 10.1091/mbc.e06-01-0089

19 De Vries, K. J. et al. Fluorescently labeled phosphatidylinositol transfer protein isoforms (alpha and beta), microinjected into fetal bovine heart endothelial cells, are targeted to distinct intracellular sites. Exp Cell Res 227, 33–39 (1996). 10.1006/excr.1996.0246

20 Ile, K. E. et al. Zebrafish class 1 phosphatidylinositol transfer proteins: PITPbeta and double cone cell outer segment integrity in retina. Traffic 11, 1151–1167 (2010). 10.1111/j.1600-0854.2010.01085.x

21 Zhao, L. et al. Individual phosphatidylinositol transfer proteins have distinct functions that do not involve lipid transfer activity. Blood Adv 7, 4233–4246 (2023). 10.1182/bloodadvances.2022008735

22 Wang, Y. J. et al. Phosphatidylinositol 4 phosphate regulates targeting of clathrin adaptor AP-1 complexes to the Golgi. Cell 114, 299–310 (2003). 10.1016/s0092-8674(03)00603-2

23 Godi, A. et al. FAPPs control Golgi-to-cell-surface membrane traffic by binding to ARF and PtdIns(4)P. Nat Cell Biol 6, 393–404 (2004). 10.1038/ncb1119

24 Capitano, M. et al. Phosphatidylinositol transfer proteins regulate megakaryocyte TGF-beta1 secretion and hematopoiesis in mice. Blood 132, 1027–1038 (2018). 10.1182/blood-2017-09-806257

25 Chambers, S. M. et al. Hematopoietic fingerprints: an expression database of stem cells and their progeny. Cell Stem Cell 1, 578–591 (2007). 10.1016/j.stem.2007.10.003

26 Di Tullio, A. et al. CCAAT/enhancer binding protein alpha (C/EBP(alpha))-induced transdifferentiation of pre-B cells into macrophages involves no overt retrodifferentiation. Proc Natl Acad Sci U S A 108, 17016–17021 (2011). 10.1073/pnas.1112169108

27 Novershtern, N. et al. Densely interconnected transcriptional circuits control cell states in human hematopoiesis. Cell 144, 296–309 (2011). 10.1016/j.cell.2011.01.004

28 Zini, R. et al. CALR mutational status identifies different disease subtypes of essential thrombocythemia showing distinct expression profiles. Blood Cancer J 7, 638 (2017). 10.1038/s41408-017-0010-2

29 Tusi, B. K. et al. Population snapshots predict early haematopoietic and erythroid hierarchies. Nature 555, 54–60 (2018). 10.1038/nature25741

30 Pietras, E. M. et al. Functionally Distinct Subsets of Lineage-Biased Multipotent Progenitors Control Blood Production in Normal and Regenerative Conditions. Cell Stem Cell 17, 35–46 (2015). 10.1016/j.stem.2015.05.003

31 Hsieh, H. H. et al. Epo-IGF1R cross talk expands stress-specific progenitors in regenerative erythropoiesis and myeloproliferative neoplasm. Blood 140, 2371–2384 (2022). 10.1182/blood.2022016741

32 Guo, G. et al. Mapping cellular hierarchy by single-cell analysis of the cell surface repertoire. Cell Stem Cell 13, 492–505 (2013). 10.1016/j.stem.2013.07.017

33 Ebner, M., Lucic, I., Leonard, T. A. & Yudushkin, I. PI(3,4,5)P(3) Engagement Restricts Akt Activity to Cellular Membranes. Mol Cell 65, 416–431 e416 (2017). 10.1016/j.molcel.2016.12.028

34 Liu, S. L. et al. Quantitative Lipid Imaging Reveals a New Signaling Function of Phosphatidylinositol-3,4-Bisphophate: Isoform- and Site-Specific Activation of Akt. Mol Cell 71, 1092–1104 e1095 (2018). 10.1016/j.molcel.2018.07.035

35 Khan, I. et al. AKT is a therapeutic target in myeloproliferative neoplasms. Leukemia 27, 1882–1890 (2013). 10.1038/leu.2013.167

36 Han, X. et al. Targeting pleckstrin-2/Akt signaling reduces proliferation in myeloproliferative neoplasm models. J Clin Invest 133 (2023). 10.1172/JCI159638

37 Davies, B. R. et al. Preclinical pharmacology of AZD5363, an inhibitor of AKT: pharmacodynamics, antitumor activity, and correlation of monotherapy activity with genetic background. Mol Cancer Ther 11, 873–887 (2012). 10.1158/1535-7163.MCT-11-0824-T

38 Turner, N. C. et al. Capivasertib in Hormone Receptor-Positive Advanced Breast Cancer. N Engl J Med 388, 2058–2070 (2023). 10.1056/NEJMoa2214131

39 Wu, H., Klingmuller, U., Besmer, P. & Lodish, H. F. Interaction of the erythropoietin and stem-cell-factor receptors. Nature 377, 242–246 (1995). 10.1038/377242a0

40 Wu, H., Klingmuller, U., Acurio, A., Hsiao, J. G. & Lodish, H. F. Functional interaction of erythropoietin and stem cell factor receptors is essential for erythroid colony formation. Proc Natl Acad Sci U S A 94, 1806–1810 (1997). 10.1073/pnas.94.5.1806

41 Zaleskas, V. M. et al. Molecular pathogenesis and therapy of polycythemia induced in mice by JAK2 V617F. PLoS One 1, e18 (2006). 10.1371/journal.pone.0000018

42 Lacout, C. et al. JAK2V617F expression in murine hematopoietic cells leads to MPD mimicking human PV with secondary myelofibrosis. Blood 108, 1652–1660 (2006). 10.1182/blood-2006-02-002030

43 Marty, C. et al. Myeloproliferative neoplasm induced by constitutive expression of JAK2V617F in knock-in mice. Blood 116, 783–787 (2010). 10.1182/blood-2009-12-257063

44 Yan, D., Hutchison, R. E. & Mohi, G. Critical requirement for Stat5 in a mouse model of polycythemia vera. Blood 119, 3539–3549 (2012). 10.1182/blood-2011-03-345215

45 Woods, B. et al. Activation of JAK/STAT Signaling in Megakaryocytes Sustains Myeloproliferation In Vivo. Clin Cancer Res 25, 5901–5912 (2019). 10.1158/1078-0432.CCR-18-4089

46 van Tiel, C. M. et al. The Golgi localization of phosphatidylinositol transfer protein beta requires the protein kinase C-dependent phosphorylation of serine 262 and is essential for maintaining plasma membrane sphingomyelin levels. J Biol Chem 277, 22447–22452 (2002). 10.1074/jbc.M201532200

47 Morgan, C. P. et al. Differential expression of a C-terminal splice variant of phosphatidylinositol transfer protein beta lacking the constitutive-phosphorylated Ser262 that localizes to the Golgi compartment. Biochem J 398, 411–421 (2006). 10.1042/BJ20060420

48 Vieira, N. M. et al. Repression of phosphatidylinositol transfer protein alpha ameliorates the pathology of Duchenne muscular dystrophy. Proc Natl Acad Sci U S A 114, 6080–6085 (2017). 10.1073/pnas.1703556114

49 Fiskus, W. et al. Dual PI3K/AKT/mTOR inhibitor BEZ235 synergistically enhances the activity of JAK2 inhibitor against cultured and primary human myeloproliferative neoplasm cells. Mol Cancer Ther 12, 577–588 (2013). 10.1158/1535-7163.MCT-12-0862

50 Morioka, S. et al. A mass spectrometric method for in-depth profiling of phosphoinositide regioisomers and their disease-associated regulation. Nat Commun 13, 83 (2022). 10.1038/s41467-021-27648-z

51 Wu, J. Z. et al. Sorting nexin 10 regulates lysosomal ionic homeostasis via ClC-7 by controlling PI(3,5)P2. J Cell Biol 224 (2025). 10.1083/jcb.202408174

52 Koizumi, A. et al. Increased fatty acyl saturation of phosphatidylinositol phosphates in prostate cancer progression. Sci Rep 9, 13257 (2019). 10.1038/s41598-019-49744-3

