## Supplemental Figures and Tables for "PITPβ Drives JAK2 V617F-Mediated Myeloproliferative Neoplasms by Promoting PtdIns(3,4)P_₂_-Dependent AKT Hyperactivation"

### Vantsev et al, Supplemental Figure 1

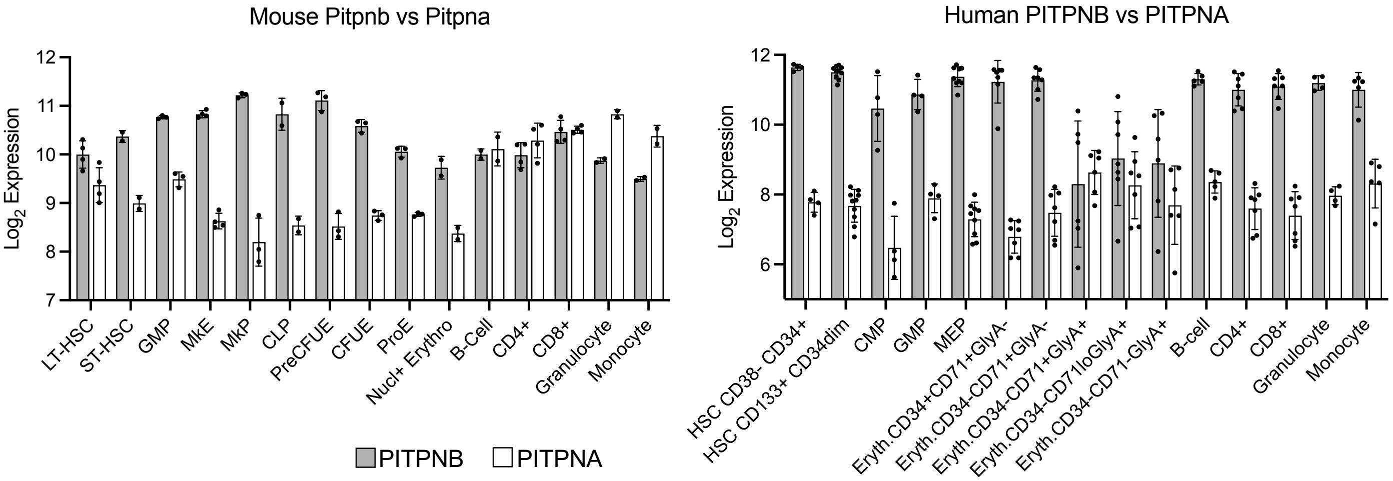

**Vantsev et al, Supplemental Figure 2**

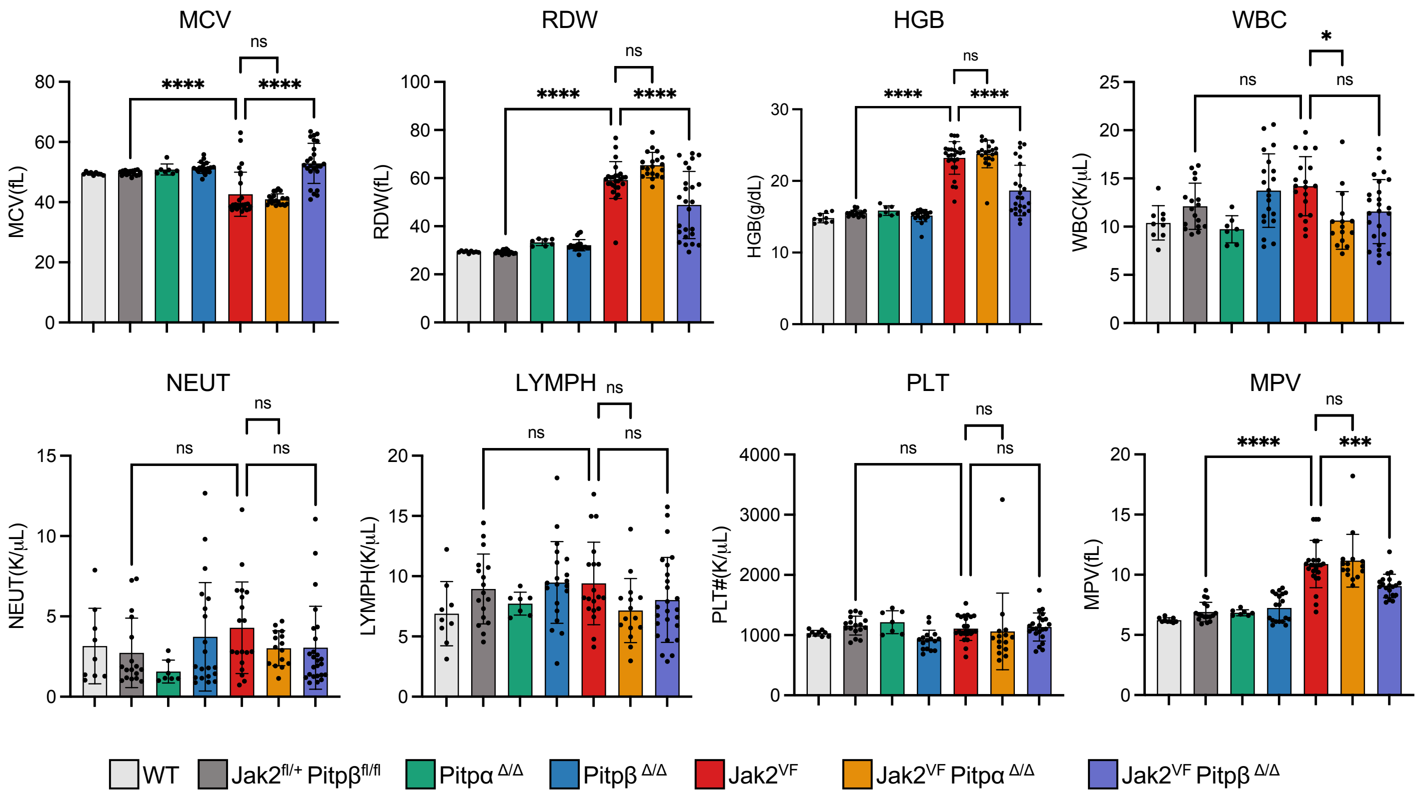

**Vantsev et al, Supplemental Figure 3**

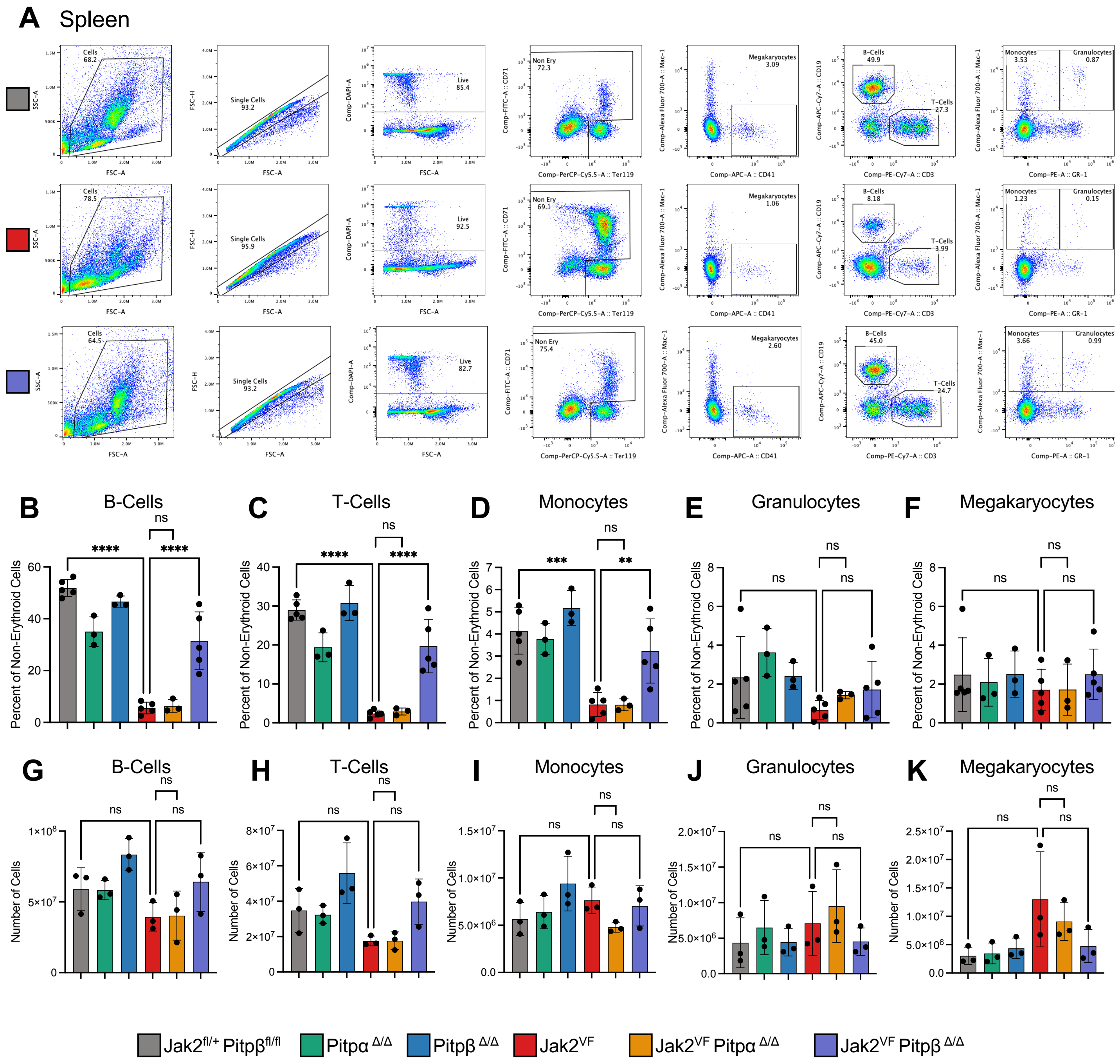

**Vantsev et al, Supplemental Figure 4**

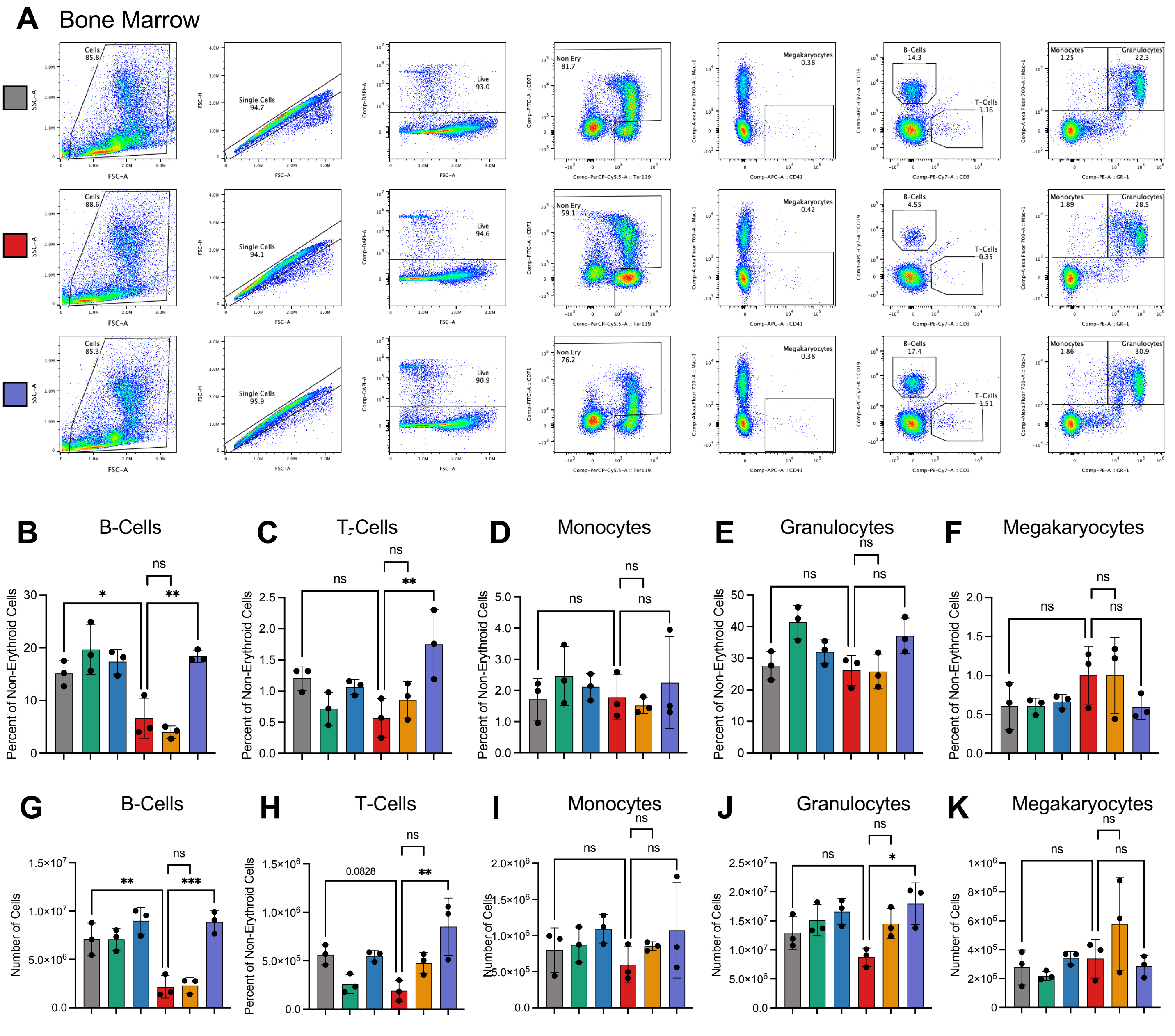


Jak2$^{fl/+}$ Pitpβ$^{fl/fl}$ | Pitpα $^{Δ/Δ}$ | Pitpβ $^{Δ/Δ}$ | Jak2$^{VF}$ | Jak2$^{VF}$ Pitpα $^{Δ/Δ}$ | Jak2$^{VF}$ Pitpβ $^{Δ/Δ}$

Vantsev et al, Supplemental Figure 5

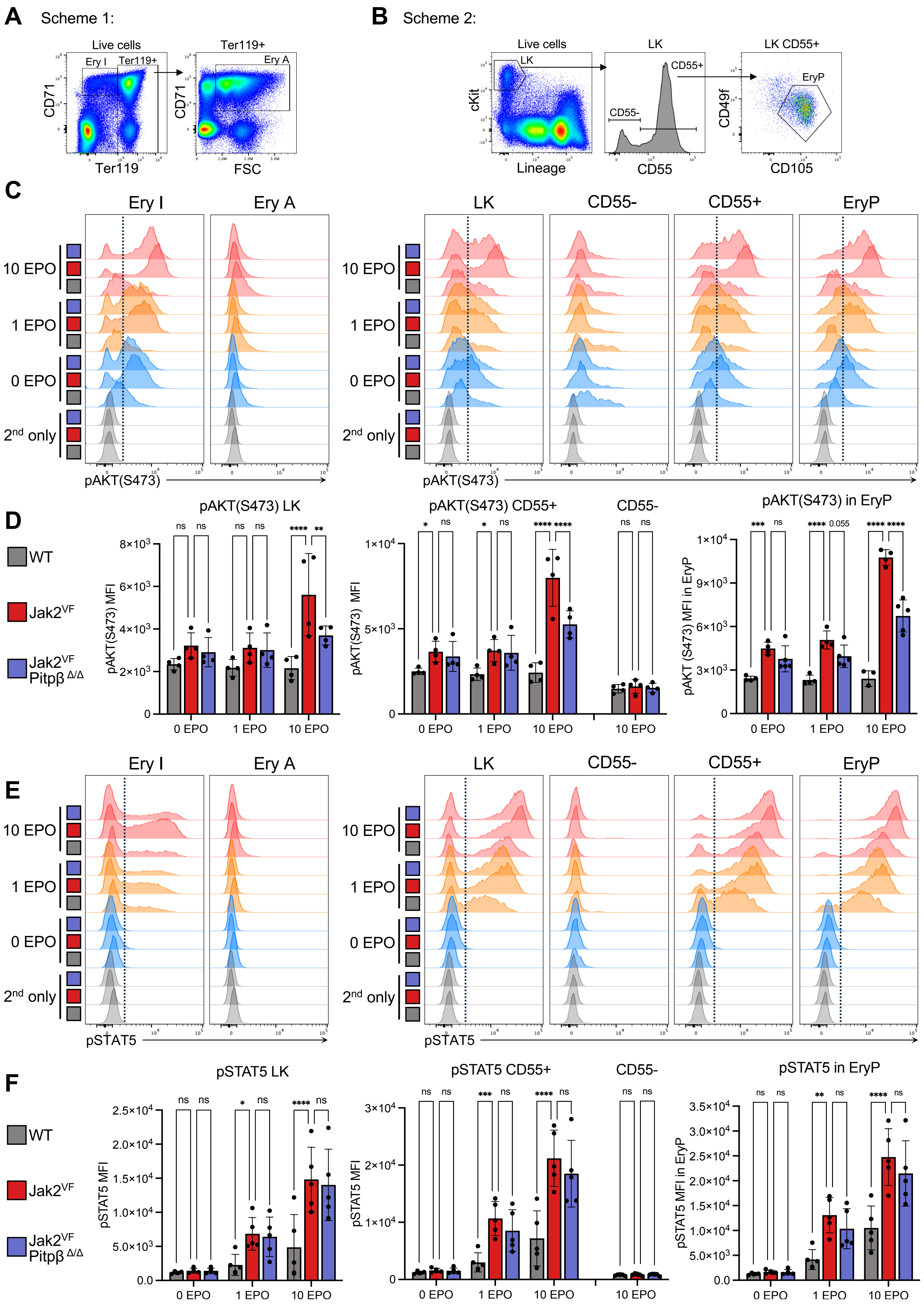

Vantsev et al, Supplemental Figure 6

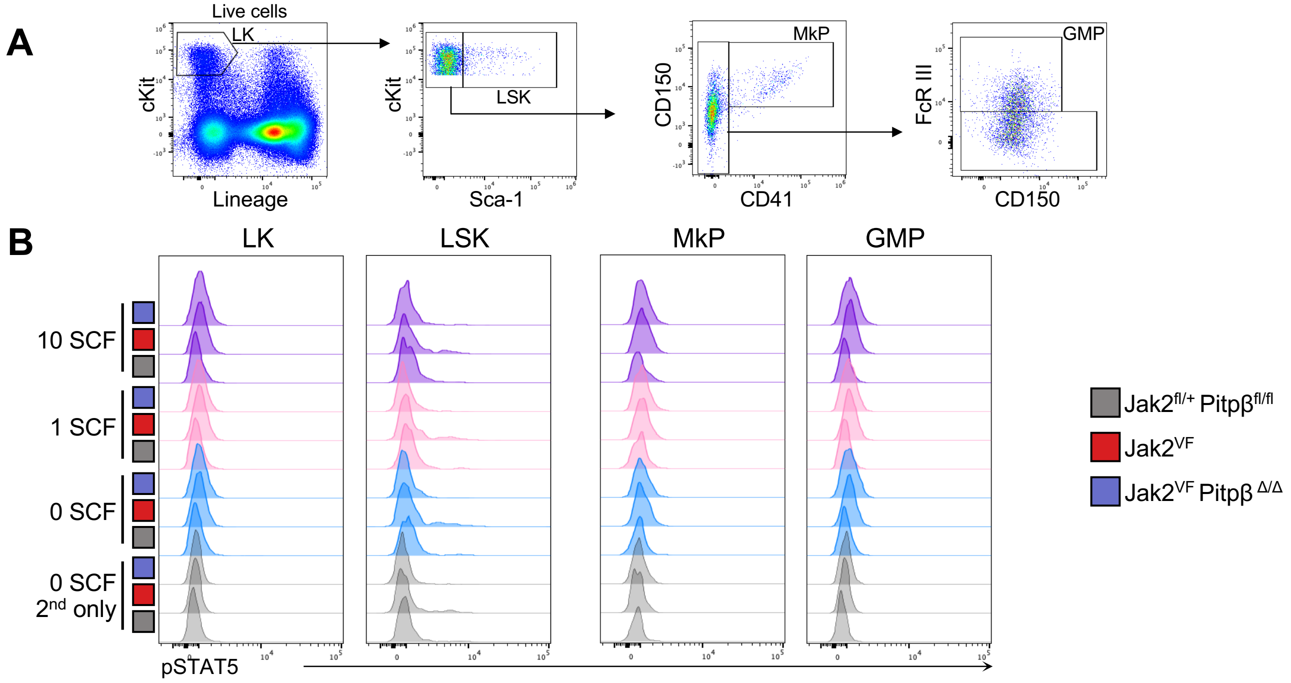

**PITPβ Drives JAK2 V617F-Mediated Myeloproliferative Neoplasms by Promoting PtdIns(3,4)$P_2$-Dependent AKT Hyperactivation**

#### SUPPLEMENTAL TABLES

##### Supplemental Table 1, Lineage Panel

| Antigen | Fluorochrome | Clone | Supplier | Identifier | Dilution |
|---|---|---|---|---|---|
| CD41 | RB613 | MWReg30 | BD | 759280 | 1:4000 |
| Gr-1 | RB705 | RB6-8C5 | BD | 570633 | 1:2000 |
| Mac-1 | APC | M1/70 | eBioscience | 17-0112-82 | 1:800 |
| Ter119 | PE | TER-119 | BD | 553673 | 1:500 |
| CD71 | FITC | C2 | BD | 553267 | 1:200 |
| CD19 | APC-Cy7 | eBio1D3 | eBioscience | 17-0193-82 | 1:200 |
| CD3 | PE-Cy7 | 145-2c11 | eBioscience | 25-0031-82 | 1:100 |
| CD45.1 | eFluor450 | A20 | eBioscience | 48-0453-82 | 1:100 |
| CD45.2 | BUV395 | 104 | BD | 564616 | 1:100 |

##### Supplemental Table 2, Lineage Biotin Panel

| Antigen | Fluorochrome | Clone | Supplier | Identifier |
|---|---|---|---|---|
| Mac-1 | Biotin | M1/70 | eBioscience | 13-0112-86 |
| Gr-1 | Biotin | RB6-8C5 | eBioscience | 13-5931-86 |
| B220 | Biotin | M1/70 | eBioscience | 13-0452-85 |
| CD19 | Biotin | 1D3 | eBioscience | 13-0193-85 |
| Ter119 | Biotin | TER-119 | eBioscience | 13-5921-85 |
| CD5 | Biotin | 53-7.3 | eBioscience | 13-0051-85 |
| CD4 | Biotin | GK1.5 | eBioscience | 13-0041-86 |
| CD8 | Biotin | 53-6.7 | eBioscience | 13-0081-85 |

**Supplemental Table 3, HSPC Panel**

| Antigen | Fluorochrome | Clone | Supplier | Identifier | Dilution |
|---|---|---|---|---|---|
| CD41 | RB613 | MWReg30 | BD | 759280 | 1:4000 |
| Streptavidin | PE-TxR | | Invitrogen | SA1017 | 1:2000 |
| FcγRIII | RB780 | 2.4G2 | BD | 569324 | 1:1000 |
| c-Kit | APC-eFlour780 | 2B8 | eBioscience | 47-1171-82 | 1:1000 |
| CD48 | AF700 | HM48-1 | Biolegend | 103426 | 1:800 |
| CD55 | AF647 | RIKO-3 | Biolegend | 131806 | 1:800 |
| Sca1 | BV650 | D7 | Biolegend | 108143 | 1:400 |
| CD105 | PE-Cy5 | MJ7/18 | Biolegend | 120427 | 1:400 |
| CD150 | BV711 | TC15-12F12.2 | Biolegend | 115941 | 1:250 |
| CD71 | FITC | C2 | BD | 553267 | 1:200 |
| CD49f | BV421 | GoH3 | Biolegend | 313623 | 1:100 |
| Flt3 | PE | A2F10.1 | BD | 553842 | 1:100 |
| CD45.1 | PE-Cy7 | A20 | BD | 560578 | 1:100 |
| CD45.2 | BUV395 | 104 | BD | 564616 | 1:100 |

**Supplemental Table 4, Sorter Panel**

| Antigen | Fluorochrome | Clone | Supplier | Identifier | Dilution |
|---|---|---|---|---|---|
| cKit | APC-eFluor 780 | 2B8 | eBioscience | 47-1171-82 | 1:200 |
| CD55 | AF647 | RIKO-3 | Biolegend | 131806 | 1:200 |
| CD105 | PE | MJ7/18 | eBioscience | 12-1051-81 | 1:200 |
| CD49f | BV421 | GoH3 | Biolegend | 313623 | 1:200 |

**Supplemental Table 5, Pflow Erythroid Panel:**

| Antigen | Fluorochrome | Clone | Supplier | Identifier | Dilution |
|---|---|---|---|---|---|
| CD105 | BUV395 | MJ7/18 | BD | 740316 | 1:200 |
| CD71 | BUV496 | C2 | BD | 741066 | 1:250 |
| CD49f | BV421 | GoH3 | Biolegend | 313623 | 1:200 |
| Sca-1 | BV650 | D7 | Biolegend | 108143 | 1:400 |
| CD150 | BV711 | TC15-12F12.2 | BioLegend | 115941 | 1:500 |
| CD3 | FITC | 145-2c11 | eBioscience | 11-0031-86 | 1:1000 |
| B220 | FITC | RA3-6B2 | eBioscience | 11-0452-82 | 1:1000 |
| Gr-1 | FITC | RB6-8C5 | BioLegend | 108406 | 1:4000 |
| Mac-1 | FITC | M1/70 | eBioscience | 11-0112-85 | 1:1000 |
| Ter-119 | RB780 | TER-119 | BD | 569366 | 1:500 |
| CD55 | Alexa Flour 647 | RIKO-3 | Biolegend | 131806 | 1:800 |
| CD48 | Alexa Flour 700 | HM48-1 | Biolegend | 103426 | 1:500 |
| cKit | StarBright Red 815 | 2B8 | Bio-Rad | MCA1365SBR815 | 1:250 |
| Anti-Rabbit | Alexa Fluor 568 | *polyclonal* | Invitrogen | A-11011 | 1:2000 |

**Supplemental Table 6, Pflow Myeloid Panel:**

| Antigen | Fluorochrome | Clone | Supplier | Identifier | Dilution |
|---|---|---|---|---|---|
| CD105 | BUV395 | MJ7/18 | BD | 740316 | 1:200 |
| CD135 | BV421 | A2F10 | Biolegend | 135314 | 1:250 |
| Sca-1 | BV650 | D7 | Biolegend | 108143 | 1:400 |
| CD150 | BV711 | TC15-12F12.2 | BioLegend | 115941 | 1:250 |
| CD3 | FITC | 145-2c11 | eBioscience | 11-0031-86 | 1:1000 |
| B220 | FITC | RA3-6B2 | eBioscience | 11-0452-82 | 1:1000 |
| Ter119 | FITC | TER-119 | eBioscience | 11-5921-81 | 1:500 |
| CD41 | RB613 | MWReg30 | BD | 759280 | 1:4000 |
| Gr-1 | RB705 | RB6-8C5 | BD | 570633 | 1:2000 |
| FcγRIII | RB780 | 2.4G2 | BD | 569324 | 1:1000 |
| Mac1 | Alexa Flour 647 | M1/70 | Biolegend | 101218 | 1:800 |
| CD48 | Alexa Flour 700 | HM48-1 | Biolegend | 103426 | 1:800 |
| cKit | StarBright Red 815 | 2B8 | Bio-Rad | MCA1365SBR815 | 1:250 |
| Anti-Rabbit | Alexa Fluor 568 | *polyclonal* | Invitrogen | A-11011 | 1:2000 |

**Supplemental Table 7, Primary phospho antibodies:**

| Antigen | Isotype | Clone | Supplier | Identifier | Dilution |
|---|---|---|---|---|---|
| p-STAT5 | Rabbit IgG | D47E7 | CST | 4322S | 1:400 |
| p-ERK1/2 | Rabbit IgG | D13.14.4E | CST | 4370S | 1:400 |
| p-AKT (S473) | Rabbit IgG | D9E | CST | 4060S | 1:400 |
| p-AKT (T308) | Rabbit IgG | D25E6 | CST | 13038S | 1:400 |

#### SUPPLEMENTAL FIGURE LEGENDS

**Supplemental Figure 1. PITPNB is more highly expressed than PITPNA in hematopoietic stem and progenitor cells in both humans and mice.** Microarray expression analysis of normal mouse[1,2] (left) and human[3] (right) hematopoietic cells. In all panels, each symbol represents a biological replicate. Bars indicate mean frequencies and error bars indicate SD.

**Supplemental Figure 2. *Pitpβ* deficiency normalizes red cell and platelet volume and partially reduces red cell distribution width.** CBC analysis of peripheral blood from WT, $Jak2^{fl/+}$ $Pitp\beta^{fl/fl}$, $Pitp\alpha^{\Delta/\Delta}$, $Pitp\beta^{\Delta/\Delta}$, $Jak2^{VF/+}$ $Pitp\alpha^{\Delta/\Delta}$, or $Jak2^{VF/+}$ $Pitp\beta^{\Delta/\Delta}$ mice; MCV: Mean Corpuscular Volume; RDW: Red Cell Distribution Width; Hemoglobin: HGB; WBC: White Blood Cell; NEUT: Neutrophil; LYMPH: Lymphocyte. PLT: Platelet; MPV: Mean Platelet Volume; In all panels, each symbol represents an individual mouse. Bars indicate mean frequencies, and error bars indicate SD. P-values were calculated using one-way ANOVA with Tukey's multiple-comparison posttests (*, p<0.05; **, p<0.01; ***, p<0.001; ****; p<0.0001).

**Supplemental Figure 3. *Pitpβ* deficiency normalizes B, T, and monocyte populations in the spleen of $Jak2^{VF}$ mice.** (A) Representative flow cytometry plots and quantification (B-K) of mature blood lineages in the spleen. (B-F) Quantification of non-Erythroid cells (excluding Ery IV: $CD71^{-}$, $Ter119^{+}$). (G-K) Quantification of the total number of non-Erythroid cells in the spleen. B-Cells: $CD19^{+}CD3^{-}$; T-Cells: $CD19^{-}CD3^{+}$; Monocytes: $Mac\text{-}1^{+}Gr\text{-}1^{-}$; Granulocytes: $Mac\text{-}1^{+}Gr\text{-}1^{+}$; Megakaryocytes: $Mac\text{-}1^{-}CD41^{+}$. In all relevant panels, each symbol represents an individual mouse. Bars indicate mean frequencies, and error bars indicate SD. P-values were calculated using one-way ANOVA with Tukey's multiple-comparison posttests (*, p<0.05; **, p<0.01; ***, p<0.001; ****; p<0.0001).

**Supplemental Figure 4. *Pitpβ* deficiency normalizes B and T-cell populations in the bone marrow of $Jak2^{VF}$ mice.** (A) Representative flow cytometry plots and quantification (B-K) of mature blood lineages in the BM. (B-F) Quantification of non-Erythroid cells (excluding Ery IV: $CD71^{-}$, $Ter119^{+}$). (G-K) Quantification of the total number of non-Erythroid cells in the BM. B-Cells: $CD19^{+}CD3^{-}$; T-Cells: $CD19^{-}CD3^{+}$; Monocytes: $Mac\text{-}1^{+}Gr\text{-}1^{-}$; Granulocytes: $Mac\text{-}1^{+}Gr\text{-}1^{+}$; Megakaryocytes: $Mac\text{-}1^{-}CD41^{+}$. In all relevant panels, each symbol represents an individual mouse. Bars indicate mean frequencies, and error bars indicate SD. P-values were calculated using one-way ANOVA with Tukey's multiple-comparison posttests (*, p<0.05; **, p<0.01; ***, p<0.001; ****; p<0.0001).

**Supplemental Figure 5. *Pitpβ* is required for EPO-mediated malignant AKT activation, but not STAT5, in all tested splenic progenitors of $Jak2^{VF}$ mice.** (A) Representative flow cytometry plots and gating strategy of fixed and permeabilized splenic cells. (C) Representative flow cytometry plots and (D) quantification of median fluorescence intensity (MFI) of pAKT(S473) in indicated populations. (E) Representative flow cytometry plots and (F) quantification of MFI of pSTAT5 in indicated populations.

Ery I: $CD71^{+}Ter119^{-}$; Ery A: $Ter119^{+}CD71^{+}$, $FSC^{high}$; LK: $Lin^{-}cKit^{+}$; EryP: LK $CD55^{+}CD105^{+}CD49f^{-}$. In all relevant panels, each symbol represents an individual mouse. Bars indicate mean frequencies, and error bars indicate SD. P-values were calculated using two-way ANOVA with Tukey's multiple-comparison posttests (*, p<0.05; **, p<0.01; ***, p<0.001; ****; p<0.0001).

**Supplemental Figure 6. SCF does not stimulate STAT5 phosphorylation.** (A) Representative flow cytometry plots and gating strategy of fixed and permeabilized splenic cells. (B) Representative flow cytometry plots of pSTAT5 MFI in indicated populations LK: $Lin^{-}cKit^{+}$; LSK: $Lin^{-}cKit^{+}Sca\text{-}1^{+}$; $LKS^{-}$: $Lin^{-}cKit^{+}Sca\text{-}1^{-}$; MkP: $LKS^{-}$ $CD150^{+}CD41^{+}$; GMP: $LKS^{-}$ $CD41^{-}Fc\gamma RIII^{+}$.

#### REFERENCES FOR SUPPLEMENTAL INFORMATION


1 Chambers, S. M. *et al.* Hematopoietic fingerprints: an expression database of stem cells and their progeny. *Cell Stem Cell* **1**, 578–591 (2007). https://doi.org/10.1016/j.stem.2007.10.003

2 Di Tullio, A. *et al.* CCAAT/enhancer binding protein alpha (C/EBP(alpha))-induced transdifferentiation of pre-B cells into macrophages involves no overt retrodifferentiation. *Proc Natl Acad Sci U S A* **108**, 17016–17021 (2011). https://doi.org/10.1073/pnas.1112169108

3 Novershtern, N. *et al.* Densely interconnected transcriptional circuits control cell states in human hematopoiesis. *Cell* **144**, 296–309 (2011). https://doi.org/10.1016/j.cell.2011.01.004
